# Single cell multi-omics analysis reveals novel roles for DNA methylation in sensory neuron injury responses

**DOI:** 10.1101/572446

**Authors:** Youjin Hu, Qin An, Guoping Fan

## Abstract

DNA methylation is implicated in neuronal injury response and regeneration, but its role in regulating stable transcription changes in different types of dorsal root ganglion (DRG) neurons is unclear. In this study, we simultaneously profiled both the DNA methylome and mRNA transcriptome from single DRG neurons at different ages under either control or peripheral nerve injury condition. We found that age-related expression changes in Notch signaling genes and methylation changes at Notch receptor binding sites are associated with the age-dependent decline in peripheral nerve regeneration potential. Moreover, selective hypomethylation of AP-1 complex binding sites on regeneration-associated gene (RAG) promoters coincides with RAG transcriptional upregulation after injury. Consistent with the findings that different subtypes of DRG neurons exhibit distinct methylome changes upon injury responses, in a hybrid CAST/Ei; C57BL/6 genetic background, we further observed allele-specific gene regulation and methylation changes for many RAGs after injury. We suggest that the genetic background determines distinct allele-specific DNA methylomes, which contribute to age-dependent regulation and neuronal subtype-specific injury-responses in different mouse strains.

## Introduction

Nerve injury in the peripheral nerve system (PNS) induces sustained injury responses including chronic pain and the activation of regeneration-associated genes (RAGs), both of which involve stable changes of gene expression (1–4). It is known that the regeneration potential of peripheral neurons declines with age as indicated by the reduction of RAG expression in aged animals. However, the molecular mechanism underlying the age-related injury responses is unclear. One of the major epigenetic regulators, DNA methylation, is associated with changes in transcriptional programs in response to intrinsic and extrinsic signals. Previous genomic studies examined the relationship between DNA methylation and transcriptional change in sensory neurons via transcriptome and methylome profiling in whole dorsal root ganglia (DRG) (5–8). However, these results are confounded by the contamination of non-neuronal cells as well as the heterogeneity of subtypes of DRG neurons. Our recent study using single cell RNA-seq showed that different types of DRG neurons exhibit distinctive transcriptional response to injury (9). Therefore, in this study, we simultaneously profiled the transcriptome and DNA methylation dynamics caused by injury in single DRG neurons at different ages (10).

## Results

### Simultaneous profiling of transcriptome and DNA methylome in individual DRG neurons

Specifically, we performed sciatic nerve transection in 1-month, 8-month and one-year old mice in a C57BL/6; CAST/Ei hybrid strain background. After 14 days post sciatic nerve transection, we manually isolated L4 and L5 DRG neurons from both contralateral (intact) and ipsilateral (injured) side DRGs and performed simultaneous single cell-methylome and transcriptome (scMT-seq) profiling (n= 83). We successfully obtained DNA methylomes from 53 single cells, each of which has 4.9 million CpGs covered on average (Fig. S1A, S1B). Importantly, among these 53 cells with methylome profiles, 35 neurons also have their transcriptome profiled, with 12,365 RNA transcripts detected on average. As expected, the analysis of the DNA methylome and transcriptome from the same cell revealed a negative correlation between expression and methylation levels around TSS (Fig. S2) (10, 11). In addition, we separately profiled either transcriptome or DNA methylome pooled in approximately 20 DRG neurons per sample using µRNA-seq (ultra-low input RNA-seq) and µWGBS (ultra-low input WGBS), respectively. Sixteen µRNA-seq samples passed quality control with 13,261 genes detected on average, and none of them have high glial and astrocyte marker gene expression (Fig. S3). Twenty-two µWGBS samples passed quality control with 14 million CpG sites covered per sample on average (Fig. S4A, S4B). With these high-quality data of either single neuron or pooled 20-neuron samples, we performed detailed bioinformatic analysis of age-dependent changes of DNA methylomes and transcriptomes at both the genome-scale and allele-specific level (Table S1–S3).

### Age-dependent changes in the DNA methylome are correlated with a reduction of transcription in Notch signaling in aged DRG neurons

To determine the age-dependent impact on the transcriptome following peripheral nerve injury, we performed principle component analysis (PCA) analysis of 16 µRNA-seq samples (Fig. 1A). While control and injury samples are clearly separated into two clusters, samples within each cluster were also grouped by their age (Fig. 1A). We then performed Weighted Gene Co-expression Network Analysis (WGCNA) in order to understand how aging influences gene expression related to regeneration at a systemic level (12, 13). We identified 48 gene modules, 2 of which were comprised of genes specifically upregulated by injury (module-trait correlation r > 0.5) (Fig. 1B, and Fig. S5). Of note, one module consisted of genes activated after injury, but their magnitude of activation was significantly attenuated in aged samples (Fig. 1C). Gene ontology analysis showed that Notch signaling (P=3.88E-05) is enriched in this group of genes. In fact, many Notch signaling related genes including *Notch1*, *Jag1*, *Hey2*, *Daam2*, *Sostdc1*, *Sfrp5* and *Apoe* were the intra-modular hubs for this gene module according to the WGCNA measurement of intra-modular gene connectivity (kME) and gene-module significance (GS) (kME > 0.8 and GS > 0.8) (Fig. S6A). Indeed, Notch signaling ligand, receptors and target genes, such as *Notch1*, *Notch2*, *Jag1*, *Hey2*, *Heyl* and *Hes1*, were all upregulated in injured DRG neurons, and their activation magnitude was decreased in aged samples (Fig. 1D and S6B). These results indicated that Notch signaling activation by injury is attenuated during aging.

**Fig. 1:**
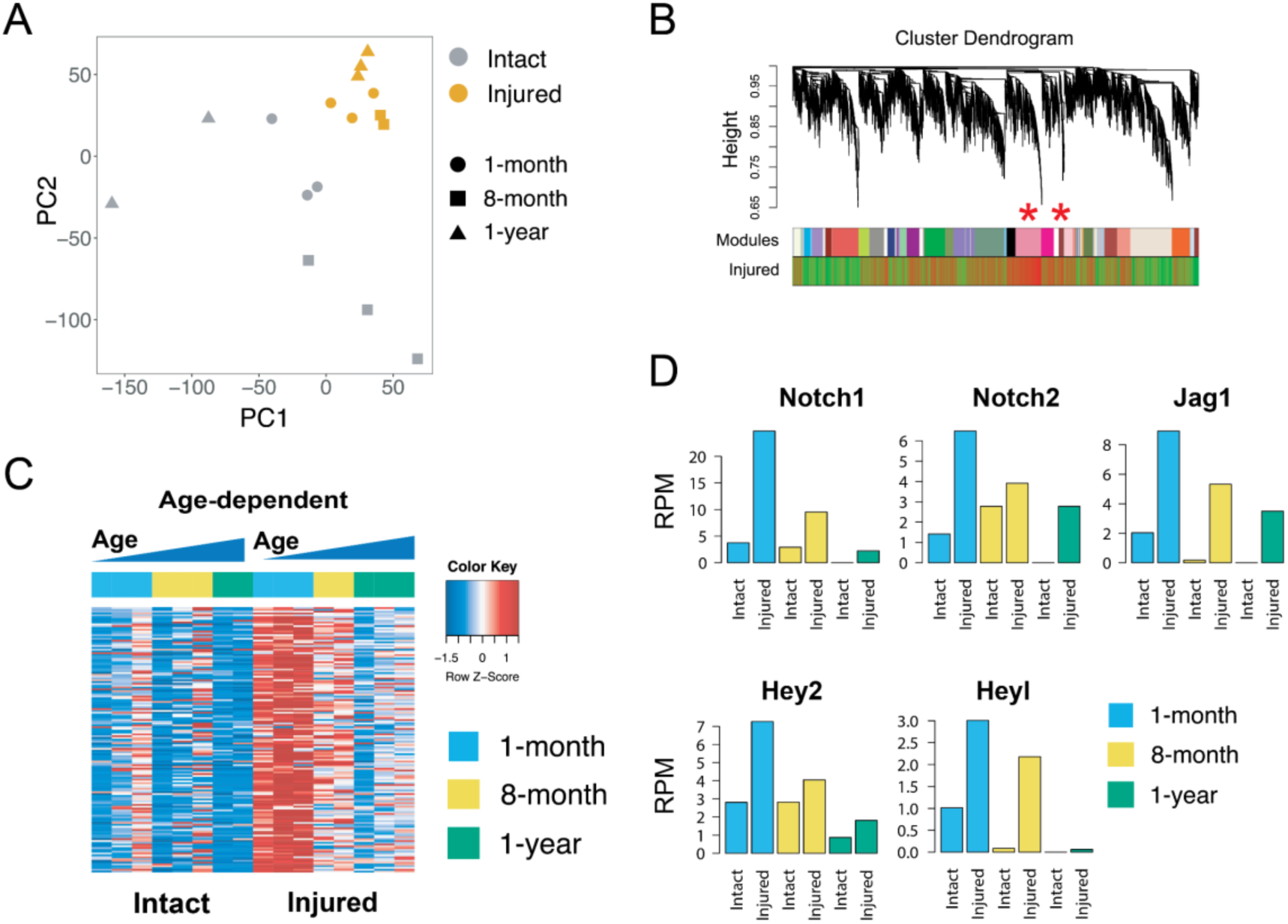
Notch signaling activation after injury is attenuated in aged neurons. **A:** Visualization of 16 µRNA-seq samples using principle component analysis showed that the samples were primarily separated by their injury condition. Within the intact or injured cluster, samples were separated by their age. **B:** WGCNA identified two gene modules (palevioletred2 and brown4, marked by red asteroids) that were specifically upregulated by injury in DRG. The hieratical clustering dendrogram represents the gene co-expression network. The “Module” and “Injured” bar underneath the dendrogram represents module assignment and correlation between sample’s injury condition of each gene on dendrogram. For the “Injured” bar, red means a gene is highly positively correlated with injury condition, and trends to be activated by injury. Green means a gene is anti-correlated with injury condition and tend to be repressed after injury. **C:** Heatmap showing the expression of genes in the brown4 (age-dependent) module. The genes in this module were activated by the injury, but the activation magnitude is attenuated in aged samples. **D:** Many notch signaling pathway components were upregulated by injury in an age-dependent manner, including Notch signaling receptors (*Notch1* and *Notch2*), ligand (*Jag1*) and downstream genes (*Hey2* and *Heyl*).

DNA methylation is involved in neuronal development, maturation and aging (14). However, the change of DNA methylation during the aging of PNS neurons has not yet been well characterized. To study this, we first analyzed the µWGBS data collected from intact neurons at different ages. Although global levels of both CpG and non-CpG methylation were similar in different age groups, we found differential DNA methylation on regulatory elements that may influence gene expression (15). Indeed, either hypermethylated (n=649) or hypomethylated (n=254) enhancers are associated with altered gene expression levels (Fig. S7C-F). Gene ontology showed differentially methylated genes were enriched in intracellular signal transduction (P=3.63E-09), central nervous system development (P=4.66E-03) and neurogenesis (P=1.45E-02). Of note, the enhancer overlap with Notch2 gene is hypermethylated during aging. These results suggest a regulatory influence of DNA methylation on gene expression including Notch signaling during the aging of PNS neurons.

DNA methylation enzymes such as Tet3 and Dnmt1 showed age-dependent upregulation after injury (Fig. S8B). We therefore examined methylome changes upon nerve injury with age. To maximize the potential of data analysis, we first combined injured or intact samples from 1-month and 1-year old mice into two pairs of meta-methylomes and identified all DMRs induced by nerve injury in each pair. We further measured the association between DMRs and the binding sites of transcription factors (TFs) using Locus Overlap Analysis (LOLA) because TF binding can trigger DNA methylation change (16). We found that for both ages, hypermethylated DMRs significantly overlap with the binding sites of CTCF and components of cohesion complex; whereas the hypomethylated DMRs were enriched with binding sites of AP-1 complex components and TFs involve in injury response. Importantly, the binding sites of AP-1 complex components were enriched in hypomethylated DMRs with a higher ratio in young neurons than older ones. Of note, the binding sites of CTCF and cohesion complex components were overlapped with hypermethylated DMRs in a higher ratio in old mice than younger ones. In addition, LOLA results indicated that the Notch1 binding sites were robustly demethylated in young mice after injury, but remained methylated in aged mice, supporting the notion that DNA methylation is involved in the attenuation of Notch signaling pathway in aged mice (Fig. S10). Given that Notch signaling activation can promote peripheral nerve regeneration (17–20), these results collectively suggest a correlation between DNA methylation changes and the suppression of Notch signaling in the reduction of regeneration potential in aged PNS neurons.

### Coordinated regulation of transcription and DNA methylation of injury-response genes in sensory neurons

The clear separation between intact and injured µRNA-seq samples in the hierarchical cluster suggests that the genetic program in sensory neurons is dramatically changed by injury (Fig. S11). We next systematically investigate how injury alters the genetic network in DRG neurons (12, 13). We identified two WGCNA modules that were comprised of genes specifically upregulated by injury. While one WGCNA module is composed of genes that exhibited age-dependent changes, the other module contains genes that are robustly activated in all injured samples (Fig. 2A, and Fig. S5). Analysis of gene ontology terms of this second module showed that genes related to neuronal regeneration including neuron projection development, axon genesis and regulation of axon extension (Fig. 2B). We identified 259 hub genes (kME > 0.8 and GS > 0.8) of this module, including many classic neural regeneration associated genes, such as *Atf3*, *Jun*, *Dcn*, *Sema6a*, *Flrt3* etc. (21–26) (Fig. 2C). These results suggested that this module represented the core regeneration-associated genetic network. Notably, among all hub genes, we found that *Sox9* had the highest centrality in this module (Fig. 2D) and is greatly induced in the injured DRG neurons (Fig. S12). Since *Sox9* plays a critical role in neurogenesis and CNS regeneration (27, 28), our study suggests a possible role for *Sox9* in injury response and regeneration of PNS neurons as well.

**Fig. 2:**
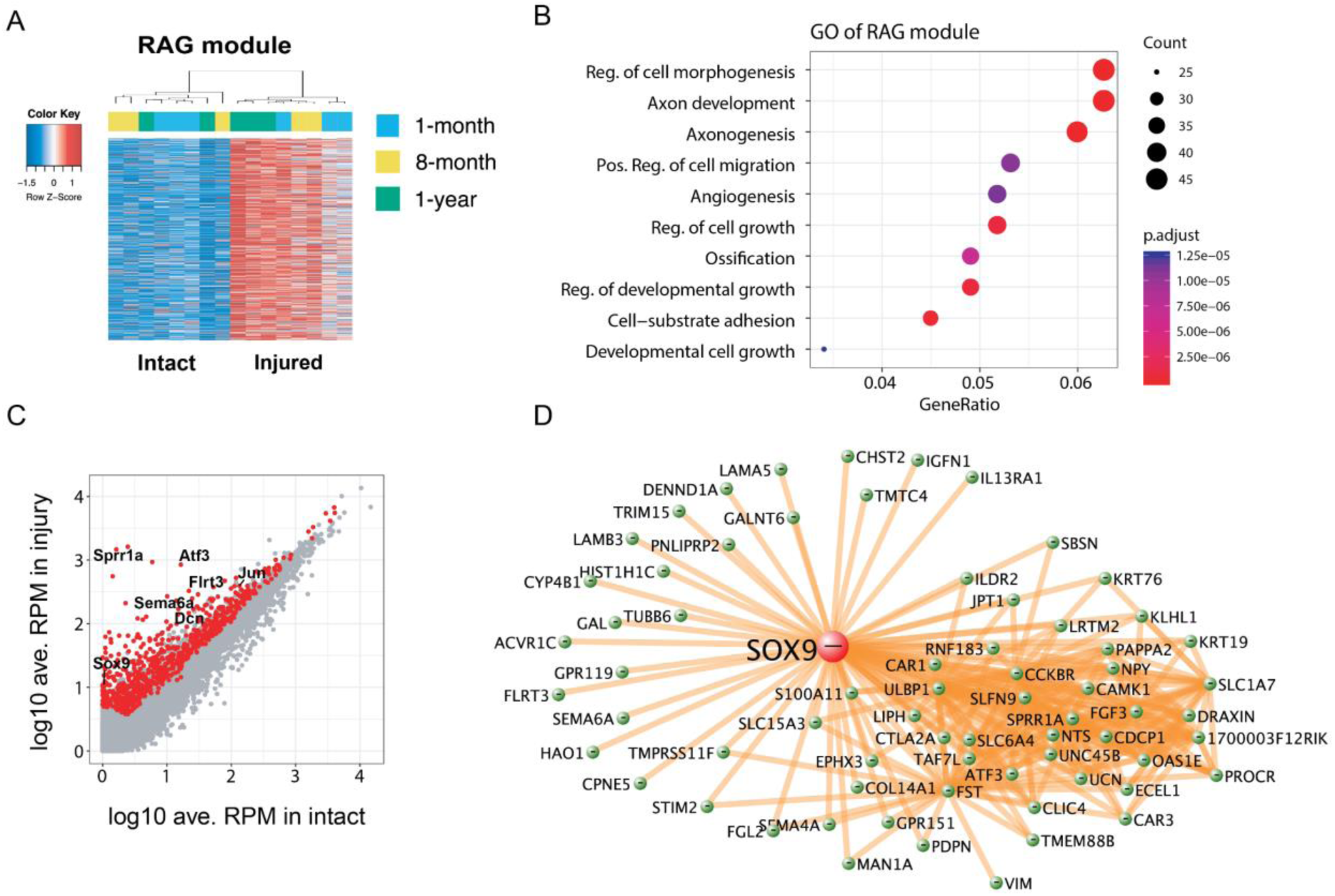
WGCNA identified a module related to neuronal regeneration. **A:** Genes in module palevioletred2 (RAG module) were expressed in a basal level in intact neurons but consistently upregulated by injury in neurons from all ages. **B:** Functional enrichment of the genes in palevioletred2 (RAG module). **C:** Scatter plot showing the average expression change of all detected genes in injured or intact samples. The genes in the palevioletred2 (RAG module, marked as red) were among the most activated genes by injury. **D:** The network structure of RAG module, showing *Sox9* is a major hub gene.

Multiple dimension reduction (MDR) and principle component analysis (PCA) of the DNA methylome data clearly separated intact and injury samples (Fig. 3A and S9C). Using a similar strategy of combining data from all injured or intact samples into two meta-DNA methylomes, we first identified all the DMRs between them. These injury-caused DMRs were enriched in regulatory elements such as CpG islands, CpG island shores and promoters that influence gene expression (Fig. S13). For example, we found that 4 TFs (*Arnt2, Atf3, Bach1* and *Sox6*) were activated and overlapped with hypomethylated regulatory elements, and two of them (*Atf3* and *Bach1*) are known to play a role in nerve regeneration (21, 22, 29, 30). Six TFs (*Camk4, Runx1, Ldb2, Myt1, Foxo3* and *Bcor*) were repressed and overlapped with hypermethylated regulatory elements, and 3 of them (*Camk4, Runx1* and *Foxo3*) involved in nerve injury response (31–33). Specifically, *Myt1* is directly bound and activated by Ascl1 (34). We found that the Ascl1 binding site on the *Myt1* promoter is hypermethylated after injury, concomitant with *Myt1* repression after injury (Fig. 3B, Fig. 3C). Overall, these results demonstrated that DNA methylation changes on regulatory elements are likely a key regulator of gene expression during injury responses of DRG neurons.

**Fig. 3:**
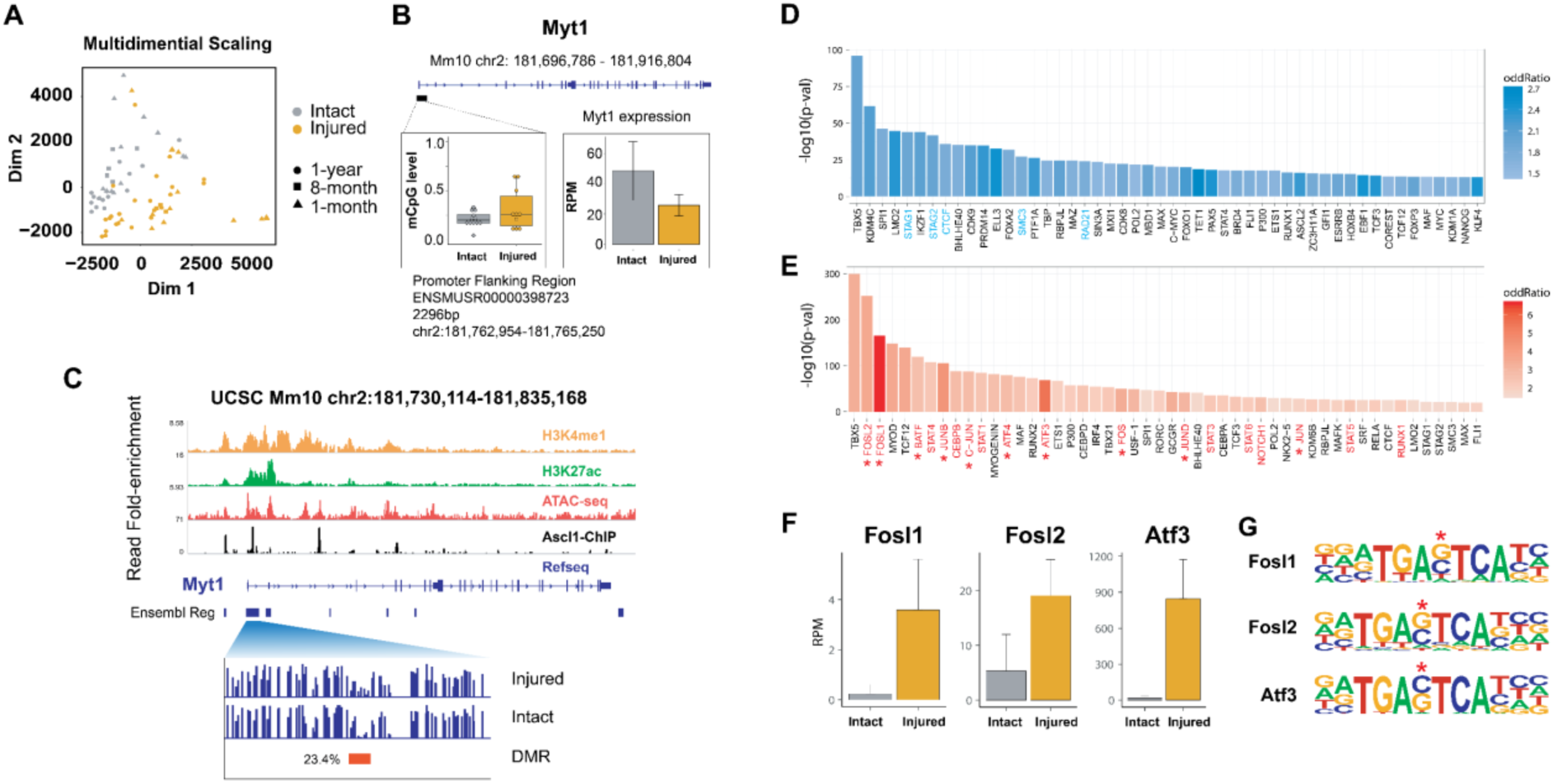
Nerve injury causes DNA methylation change in DRG neurons. **A:** Visualization of scWGBS and µWGBS data using multiple dimensional scaling (MDS). The samples were separated by their injury condition (indicated by different colors). Within intact or injured clusters, the samples were ordered by their age. **B:** A regulatory element at *Myt1* promoter is significantly hypermethylated (left panel), and *Myt1* expression is downregulated after injury (right panel). **C:** The hypermethylated regulatory element overlaps with H3K4me1, H3K27ac and Ascl1 ChIP-seq peaks, and ATAC-seq open chromatin regions. **D:** Barplot showing the significance of enrichment and enrichment fold of the association between injury-induced hyper-DMRs and transcriptional factors binding sites. Factors related to chromatin organization is highlighted as blue. **E:** Barplot showing the association with injury-induced hypo-DMRs. TFs known to involve in injury response is highlighted as red. Components of AP-1 complex components were marked by asterisks. **F:** Expression of AP-1 complex components. **G:** AP-1 binding motifs were enriched in promoter of genes in RAG module. CpG sites were marked by red asteroids.

We then examined the correlation of TF binding sites with DMRs by LOLA. For the hypermethylated DMRs, significant overlap was found with the binding sites of CTCF and components of cohesion complex (Fig. 3D). For the hypomethylated DMRs, highly significant overlap was found with binding sites of AP-1 complex components, as well as other TFs that involve nerve injury response (Fig. 3E) (35–40). It is known that AP-1 components are robustly induced in sensory neurons after injury, and indispensable for a successful axonal regeneration (21–23, 30, 41). In our datasets, the transcription of AP-1 complex components, such as *Fosl1, Fosl2, Atf3* and *Jun*, were greatly upregulated upon injury (Fig. 3F), and their binding motifs were significantly enriched in promoters of genes in this RAG module (Fig. 3G and S15). Together, these results suggest AP-1 as a common upstream regulator to activate the genes in the core regeneration-associated module after injury.

### Different subtypes of DRG neurons exhibit distinctive DNA methylome response to injury

Nerve injury in the PNS induces distinctive transcriptional response in different subtypes of DRG neurons (9). However, it is unclear whether DNA methylation is responsible for this heterogeneity in different neuronal subtypes. In our dataset, we successfully classified 35 single DRG neurons profiled by scMT-seq into 3 major subtypes based on their transcriptome, namely nonpeptidergic nociceptors (NP) (n=6), peptidergic nociceptors (PEP) (n=19) and neurofilament containing sensory neurons (NF) (n=10) (Fig. 4A). Importantly, DNA methylomes in these single DRG neurons were also simultaneously profiled, allowing us to examine the role of methylation in regulating gene expression at single-base resolution. While no significant change of global CpG methylation was found after injury in all three subtypes, non-CpG methylation is significantly increased after injury in the NF neurons (Fig. S16A) (Wilcoxon Rank Sum and Signed Rank Tests, p-value < 0.05).

**Fig 4.**
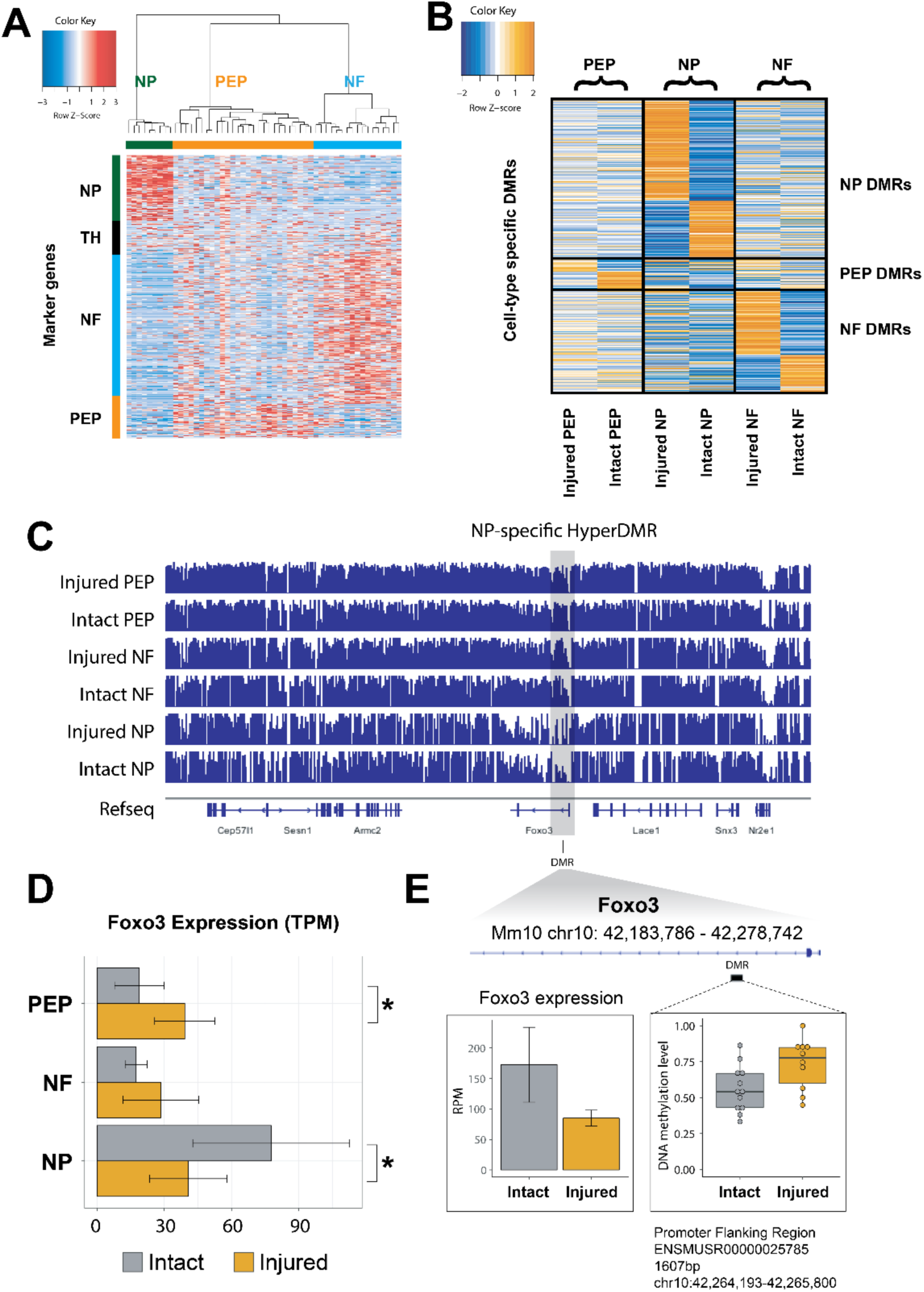
Neuron subtype-specific methylation change revealed by scMT-seq. **A:** Heatmap showing the expression of DRG neuron subtype marker genes among all DRG neurons with scRNA-seq data available, including 35 scMT-seq samples. **B:** Heatmap showing the methylation level of injury-induced DMRs identified from three neuronal subtypes separately. **C:** IGV view of the DNA methylation levels on the region around *Foxo3* gene in three different neuron subtypes. The grey box highlights the position of a NP neuron specific hypomethylated region. **D:** The expression level of *Foxo3* in three different DRG neuron subtypes. **E:** The position of the NP-specific hypermethylated element relative to *Foxo3* TSS.

To dissect the locus-specific CpG DNA methylation change, we then identified CpG DMRs caused by injury in all three types of neurons separately. We identified 225 hyper and 128 hypo DMRs in NP neurons, 28 hyper and 39 hypo DMRs in PEP neurons, and 144 hyper and 85 hypo DMRs from PEP neurons (Fig. 4B). Only a few DMRs were differentially methylated in all three types of neurons, and most of DMRs showed different methylation level change in different neuron subtypes. These results suggest that different DRG neuron subtypes exhibit distinctive DNA methylation response to injury. Notably, in NP neurons, for those genes in which their promoters exhibiting hypermethylation and down-regulated upon nerve injury, they were functionally enriched in neurogenesis, implicating their role in nerve regeneration (9). For instance, the *Foxo3* promoter is specifically hypermethylated in NP neurons but not in other neuronal cell types (Fig. 4C and Fig. 4E), and *Foxo3* is specifically repressed after injury only in NP neurons (Fig. 4D). This suggest a causal relationship between cell-type specific methylation and transcriptional changes. Taken together, our results revealed that the distinctive DNA methylation patterns in subtype neurons contribute to different injury responses in different DRG neurons.

### Integrative analysis of allele-specific DNA methylation and gene expression suggests a methylation-mediated cis-regulatory mechanism underlying the enhanced regeneration potential of CAST/Ei mice

Previous studies showed that CAST/Ei mice exhibit an enhanced axonal regeneration ability compared to C57BL/6 mice due to strain-specific regulation of a subset of regeneration-associated genes (RAGs) (42). However, it is unclear whether the regulatory mechanism is trans or cis on the expression of these RAGs (42). In our F1 mice of C57BL/6 × CAST/Ei hybrid strain, we have an opportunity to determine the expression ratio between two alleles and deduce cis- or trans-regulatory effects upon injury. We found that many genes showed unbalanced allelic expression and coordinated unbalanced promoter allelic methylation. Specifically, under intact condition, 668 genes showed higher expression from CAST/Ei allele, and 138 of them were hypomethylated on promoter at CAST/Ei allele. Similarly, 976 genes showed higher expression from C57BL/6 allele, and 231 of them were hypomethylated on promoter at C57BL/6 allele. From all 369 genes that showed unbalanced allelic expression and coordinated unbalanced promoter allelic methylation, 14 were known imprinted genes. The significant overlap (Fisher exact test, p-value < 0.05, after excluding imprinting genes and sexual chromosome associated genes) between unbalanced allelic expressed and methylated genes suggested that genetic background could shape local DNA methylation, which then influence nearby gene expression.

We then identified genes whose two alleles were regulated differentially after injury. We found 782 genes in which the CAST/Ei to C57BL/6 allele expression ratio was significantly increased, and vice versa, 410 genes in which CAST/Ei to C57BL/6 allele ratio was significantly decreased after injury (Fisher exact test, BH corrected p-value < 0.05, delta ratio change > 0.2) (Fig. 5A). All these two group genes contains 10% more strain-specific SNPs on their promoters, and were significantly enriched in CAST/Ei strain-specific RAGs (Hypergeometric test, p-value = 0.0245) (42). For example, *Inhba* and *Axl* were the top 1^st^ and 3^rd^ CAST/Ei-specific RAGs that strongly correlated with DRG regeneration ability, and hyper-activation of *Inhba* in CAST/Ei mice DRGs after injury likely confers enhanced neuronal regeneration ability (42). Our data showed a significant upregulation of *Inhba* and *Axl* from CAST/Ei allele but not C57BL/6 allele. Similarly, Galanin (*Gal*) can promote DRG neuron regeneration capacity after injury (43), and its activation level is much higher in CAST/Ei mice than in C57BL/6 (Fig. 5B). Allelic expression analysis showed that the CAST/Ei but not C57BL/6 allele of *Gal* is greatly induced after injury, resulting in a significant increase of CAST/Ei percentage of *Gal* after injury (Fig. 5C). Our results suggest that many differentially up-regulated CAST/Ei allelic RAGs are mediated by cis-regulatory elements in CAST/Ei genes.

**Fig. 5:**
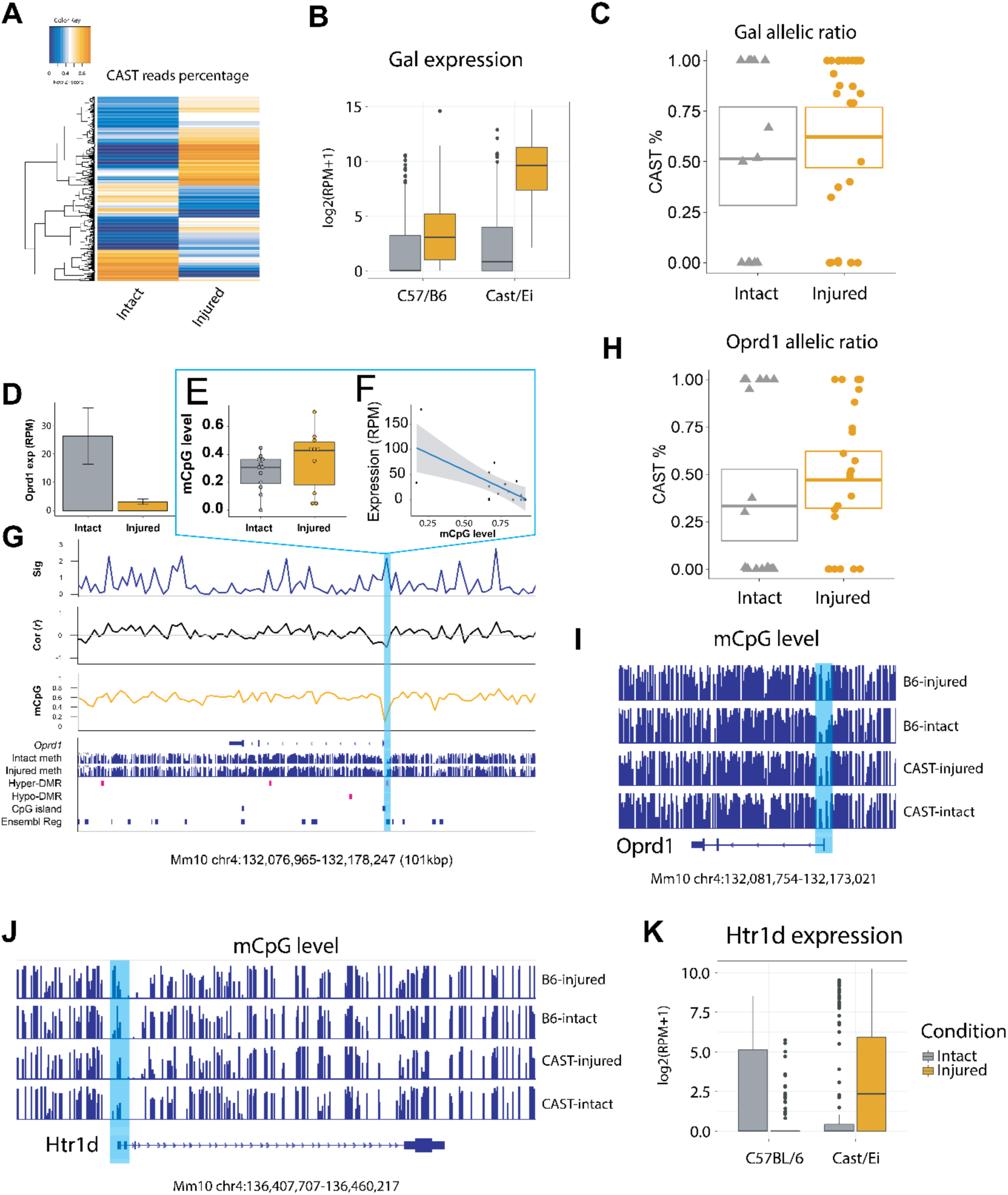
Nerve injury induces differential allelic expression in DRG neurons. **A:** Heatmap showing the gene whose allelic expression ratios significantly change upon injury. **B:** Expression change of *Gal* by injury in pure C57BL/6 and CAST/Ei mice. **C:** The boxplot showing the percentage of reads transcribed from Gal gene on CAST/Ei allele in intact and injured neurons. **D:** *Oprd1* expression is downregulated in DRG neurons by injury. **E-G**. Correlation analysis using scMT-seq data identified a differentially methylated regulatory element located at the *Oprd1* TSS. This element is hypomethylated after injury (**E**), and its methylation level is negatively correlated with *Oprd1* expression (**F**). **H:** The boxplot showing the percentage of reads transcribed from *Oprd1* gene on CAST/Ei allele in intact and injured neurons. **I:** IGV view showing the allelic DNA methylation level around *Oprd1* gene. The DMR is significantly methylated by injury on C57BL/6 allele, but not CAST/Ei allele. **J:** IGV view showing allelic DNA methylation levels around the *Hrt1d* gene. The DMR is significantly methylated by injury on C57BL/6 allele, but not CAST/Ei allele. **K:** Expression change of *Htr1d* by injury in pure C57BL/6 and CAST/Ei mice.

To determine whether local DNA methylation is one of the cis-regulatory mechanisms underlying allele-specific gene expression, we examined if methylation changes on two alleles are differentially regulated at the genome-scale upon nerve injury. Our analysis showed collectively 8,950 and 8,947 significant hyper and hypo methylated regions on C57BL/6 alleles, and 7,572 and 8,407 hyper and hypo methylated regions on CAST/Ei alleles in all DRG neurons. Among genes showing methylation changes, many were previously identified as injury-responsive genes. For example, repression of *Oprd1* in injured DRG neurons by epigenetic mechanism is responsible for chronic pain development (44). Correlation analysis identified a methylation-eQTL (meth-eQTL) near *Oprd1* transcription start site (TSS) in which DNA methylation is inversely correlated with *Oprd1* expression (Fig. 5D - 5G). Allelic expression analysis showed that *Oprd1* expression from C57BL/6 allele is significantly downregulated (Fig. 5H, Fig. S17A) together with the meth-eQTL being hypermethylated on C57BL/6 allele, indicating a differential regulation of this meth-eQTL between two alleles (Fig. 5I). *Htr1d* gene encodes 5-HT5D receptor that relates to chronic pain development after injury (45). We found that down-regulation of *Htr1d* expression from C57BL/6 allele and *Htr1d* promoter hypermethylation on C57BL/6 allele, and up-regulation of *Htr1d* expression from CAST/Ei allele and *Htr1d* promoter hypomethylation on CAST/Ei allele (Fig. 5J, Fig. 5K, Fig. S17B). Taken together, these results suggest that differential allelic methylation is one of the key cis-regulatory mechanisms underlying differential allelic expression in DRG neurons after injury (46).

## Discussion

Our study provides a comprehensive analysis of both transcriptome and methylation regulation in single DRG neurons upon injury. One of the key findings is the attenuation of Notch signaling in aged mice that contributes to the age-dependent decline in injury responses and regeneration. We found that binding sites of Notch signaling receptor tends to be hypomethylated in young DRG neurons (1-month old) compared to mature neurons at the age of one-year old. We suggest that age-dependent DNA methylation change reduces injury response and regeneration potential in old PNS neurons by the suppression of the Notch signaling (Fig. S18).

Among the enzymes responsible for DNA methylation, we found that Dnmt1 is robustly upregulated after injury across multiple datasets, regardless of age and genetic background, suggesting a potential regulatory role for Dnmt1 in PNS neurons after injury (47). Recently, *de novo* methylation activity for Dnmt1 was uncovered in Stella-mediated protection of oocyte-specific methylation (48). Our previous study has demonstrated a synergistic activity for both Dnmt1 and Dnmt3a in regulation DNA methylation in CNS neurons (49). It would be of interests to determine whether both Dnmt1 and Dnmt3a also exert its *de novo* activity in regulating methylation in postmitotic neurons in peripheral neurons upon injury.

In this study, we observed regional hypermethylation at the genome-scale in an allele-specific manner in F1 C57BL/6; CAST/Ei mice after injury. Quite a few of hyper DMRs were significantly associated with binding sites of DNA methylation sensitive chromatin structure regulators CTCF, suggesting chromatin structure in DRG neurons are robustly remodeled during injury response. Significantly, we found that allele-specific methylation patterns can serve as a cis-eQTL trait in gene regulation upon injury responses. Our finding underscores the importance of genetic background that determines allele-specific methylation in the control of regeneration potentials. Future studies shall address how genetic background, presumably dictated by species-specific DNA sequences, regulates both cell-type specific and allelic-specific DNA methylome in neuronal subtype upon nerve injury.

## Acknowledgements

We thank Dr. Matteo Pellegrini (UCLA) for his critical reading of the manuscript.

## Author contribution

G.F., Y.H. and Q.A. conceptualized the study. Y.H. performed the animal and sequencing experiments. Q.A. and G.F. performed the formal analysis and wrote the manuscript in discussion with all authors. All authors reviewed and revised the manuscript.

## Data and materials availability

All sequencing data generated in this study are available on Gene Expression Omnibus (GEO) with accession number GSE124728. All intermediate data, results and code used in the analysis are available upon reasonable request to the authors.

## Supplementary Materials

### Materials and Methods

#### Construction of DRG axotomy model and isolation DRG neurons

Mice were kept in cages under 12h light-dark condition. In this study, we used a hybrid mouse strain by crossing C57BL/6 and CAST/Ei mice, which enabled us distinguishing reads from two different alleles. Mouse DRG axotomy models were constructed following our previous published protocol (1, 2). Briefly, mice at each age group (1-month, 8-month and 1-year old, 3 mice for each age) were anesthetized by intraperitoneal injections using a mixture of ketamine (100 mg/kg) and xylazine (10 mg/kg). For each mouse, the sciatic nerve at the right hind limb was exposed at the mid-thigh level and axotomized distally, and the left side was left intact as control. The wound was sutured in two layers, and the animals were allowed to recover for 14 days. The mice were sacrificed 14 days after the axotomy surgery and lumber DRGs (L4, L5) were dissected on ice, and dissociated into single cells according to previous published protocols (3). Dissociated cells were temporarily kept on ice in DMEM medium containing 10% FBS before use.

#### µRNA-seq and µWGBS

Single DRG neurons were picked under microscope using micro-capillary pipetting. Twenty neurons were pooled into 1 µ-sample. For µRNA-seq, libraries were constructed following standard SMART-seq2 protocol. Briefly, cDNA was reverse-transcribed using Superscript III and amplified by PCR with 18 cycles. cDNA was first diluted to 0.1ng/uL, and 1uL of diluted cDNA was input for tagmentation and library amplification with 12 PCR cycles, according to standard SMART-seq2 procedure. µWGBS libraries were constructed following previous published single cell WGBS protocol with minor modification (4). Briefly, BS conversion is firstly carried out directly on the cell lysate, and single stranded DNA after BS conversion was preamplified with random primers for 5 times. After purification, another round of PCR for 12 cycles based on random priming was performed. Library quality was assessed using Tapestation assay and PAGE gel electrophoresis. The libraries were sequenced on a Hiseq 4000 machine in pair-end, 150bp mode. For 20-cell RNA-seq and 20-cell WGBS libraries, 20 cells were pooled into one tube and treated as one sample in the following library construction steps.

#### Method of single cell methylome and transcriptome sequencing (scMT-seq)

scMT-seq consists of three steps: isolation of nucleus and cytoplasma, then single cell RNA-seq using cytoplasma, and single cell whole genome bisulfate sequencing using the nucleus. Briefly, cell membrane-specific lysis buffer was used to lyse cell membrane (2). Cell nucleus was manually picked out and put into nucleus lysis buffer. The remaining cell plasma were used for transcriptomic profiling by SMART-seq2 as described previously. For the single cell whole genome bisulfate sequencing, the library was constructed by performing the single cell whole genome bisulfate sequencing protocol on the isolated nucleus (4) as described previously. Both RNA-seq and WGBS libraries were sequenced on a Hiseq 4000 machine in pair-end, 150bp mode.

#### RNA-seq data analysis

RNA-seq reads were first trimmed using TrimGalore with default parameters to remove adapter sequence and low-quality bases. Trimmed reads were then aligned to SNP-masked mouse reference genome mm10 using STAR aligner. Only reads that uniquely mapped to mm10 were retained and read count on each gene in each sample was computed using featureCounts. SNP-masked genome was generated by masking C57BL/6 and CAST/Ei strain-specific SNPs as “N” in mm10 genome using SNPSplit, to avoid potential mapping bias. Libraries with low number of uniquely mapped reads (less than 2 million reads), or libraries with low median FPKM values were excluded. Differentially expressed gene analysis was performed using SCDE. For WGCNA analysis, genes with low expression analysis (average FPKM < 1) were excluded in the subsequent analysis. WGCNA was performed on normalized gene expression level RPM (read count per million). The WGCNA co-expression network was visualized using VisAnt and in-house R scripts. Gene ontology analysis was performed using online tool: http://www.geneontology.org. The promoter transcriptional factor binding motif enrichment analysis was performed using HOMER.

#### Whole genome bisulfate sequencing data analysis

Whole genome bisulfate sequencing reads were first trimmed using TrimGalore to remove adapter and low-quality bases. The trimmed reads were then aligned to SNP-masked mm10 and CpG and non-CpG methylation signal was extracted using Bismark. Aligning reads to the masked genome can eliminate mapping bias and false-positive methylation result from C/G SNPs. Differentially methylated regions analysis was performed using R package methylkit. Locus overlap analysis was performed using R package LOLA with mm10 reference dataset.

#### Correlation analysis of DNA methylation and gene expression with scMT-seq dataset

The scMT-seq correlation analysis was performed between the expression value of a given gene A and the methylation levels among a region B. In this study, we chose gene A coding region plus the 100kbp flanking regions on both ends as B. The expression values of gene A among n cells can be represented as a column vector A_Exp_ containing n elements.

We then tile the region B into 3kbp-long, non-overlapping bins, and determined the methylation level for each bin following a Bayesian distribution with a beta distribution prior, as described before (4). By doing this procedure for each cell, we will construct a methylation value matrix B_Meth_ with m columns (each column representing one bin in region B), and n rows (each row representing one sample). Spearman correlations between A_Exp_ and each column in matrix B_Meth_ were computed. The bin whose methylation level was highly correlated with gene expression level was likely to be the cis methylation regulatory elements (cis-MREs) of gene A. The scMT-seq correlation analysis was implemented as customized R scripts.

#### Allelic expression and methylation analysis

To quantify allelic expression ratio for each gene, the aligned RNA-seq reads were first assigned to C57BL/6J or CAST/Ei allele by the strain-specific SNPs they contained using SNPsplit. The strain-specific SNP lists were downloaded from the Mouse Genomes Project (https://www.sanger.ac.uk/science/data/mouse-genomes-project). After splitting the aligned RNA-seq reads, featureCounts was used to count strain-specific reads and the expression ratio between C57BL/6J and CAST/Ei allele was determined as the ratio between C57BL/6J and CAST/Ei specific read number. To determine the allelic methylation level, the aligned WGBS reads were split using SNPsplit in BS-seq mode. Bismark was run on the split BAM files to extract the methylation signal for each allele.

---

**Fig. S1.**
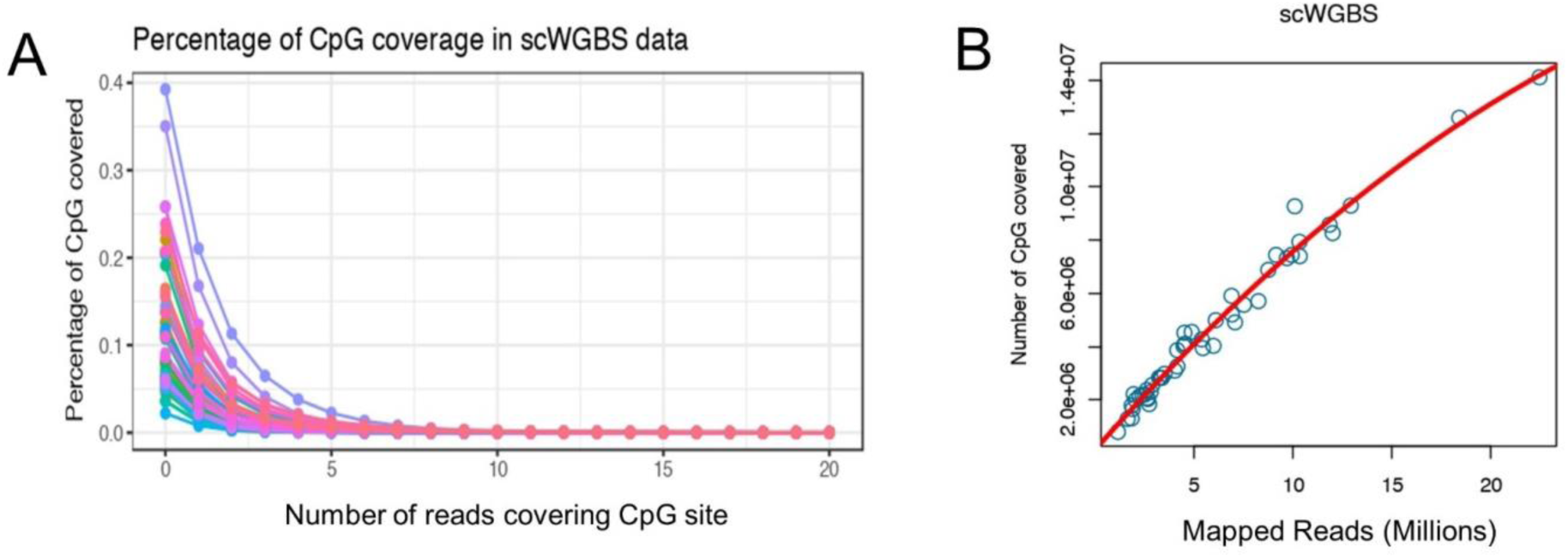
Mapping and coverage of single cell bisulfite sequencing data. **A:** The percentage of CpG sites detected in each scWGBS sample, versus the read coverage of the site. **B:** The number of CpGs covered in each sample, versus the number of mapped reads.

**Fig. S2.**
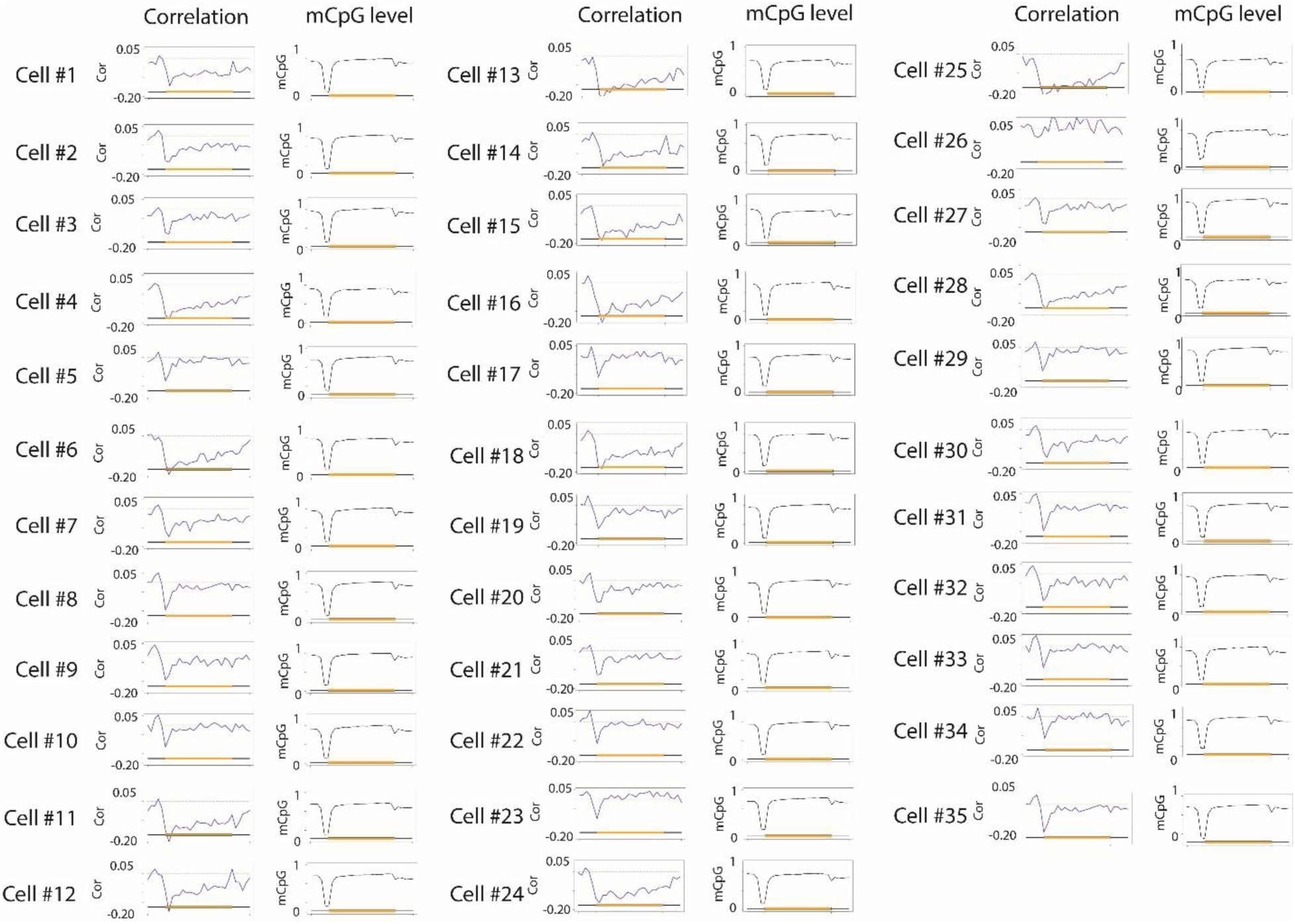
Metablots of DNA methylation and its correlation with RNA expression in a single cell. The distribution of DNA methylation across gene bodies (right panel, black line plot showing mCpG level), and spearman correlation between CpG methylation level and gene expression (left panel: correlation) in the same cell. We segmented genes from 5’ to 3’ into tiles (the gene structure is shown on the x-axis), and calculated their individual DNA methylation level and spearman correlation between their DNA methylation level and gene expression in the same segment. We found a hypomethylation valley around the TSS regions, and moderate hypermethylation on gene bodies. The correlation analysis revealed a negative correlation between promoter DNA methylation level and gene expression.

**Fig. S3.**
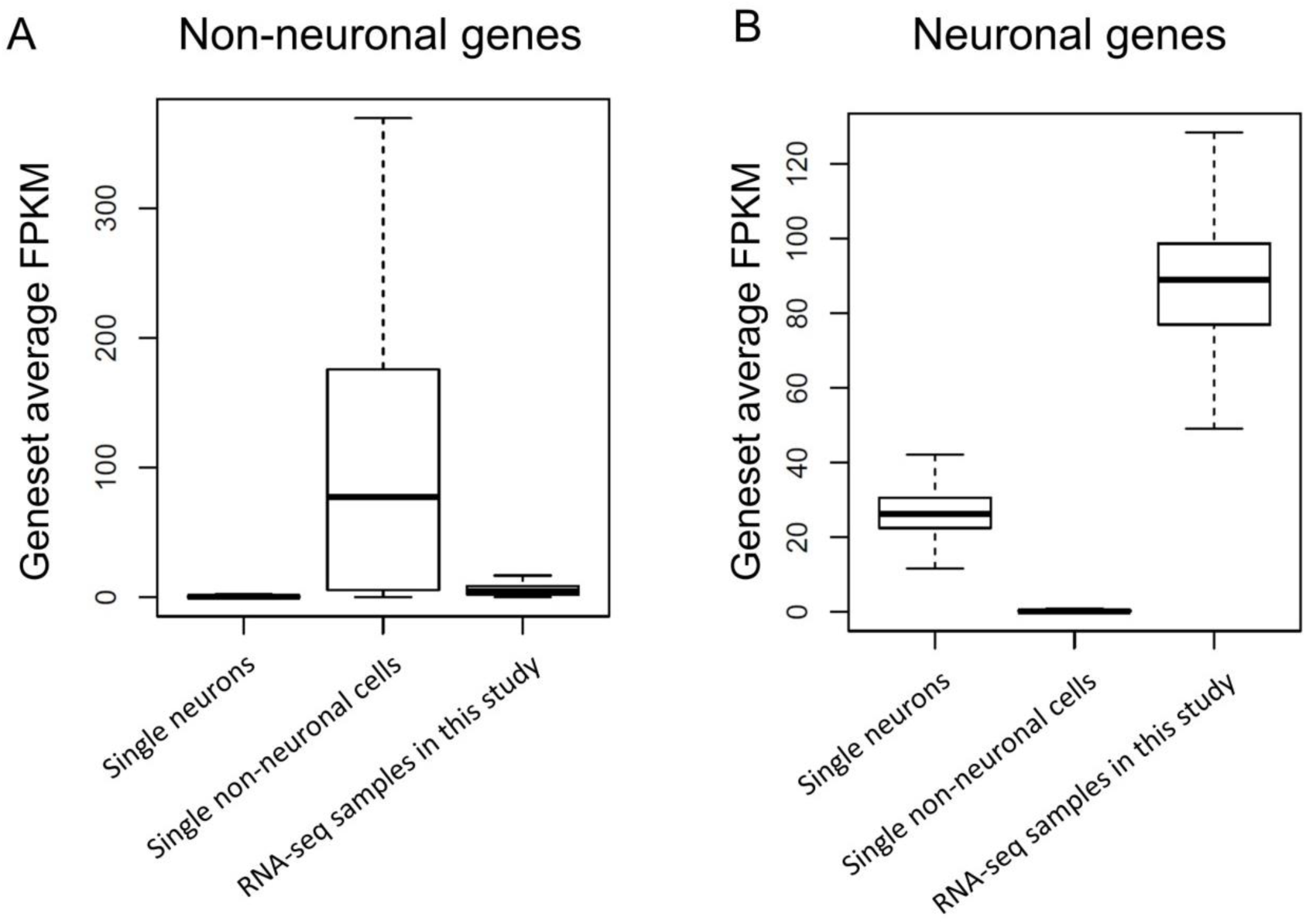
Expression pattern of neuronal specific genes and non-neuronal genes showing minimal contamination of non-neuronal cells in our DRG neuron samples. FPKM of DRG neuron-specific genes and non-neuronal genes in single DRG neurons (from Usoskin et. al (5)), single non-neuronal cells (from Usoskin et. al (5)) and µRNA-seq and scRNA-seq samples generated in this study. **A:** We found that our DRG neuron RNA-seq samples that passed quality control have minimal expression of non-neuronal marker genes. **B:** Our DRG neurons RNA-seq samples also exhibit high expression of neuronal marker genes. These results suggest minimal contamination of non-neuronal cells in our DRG neuron samples.

**Fig. S4.**
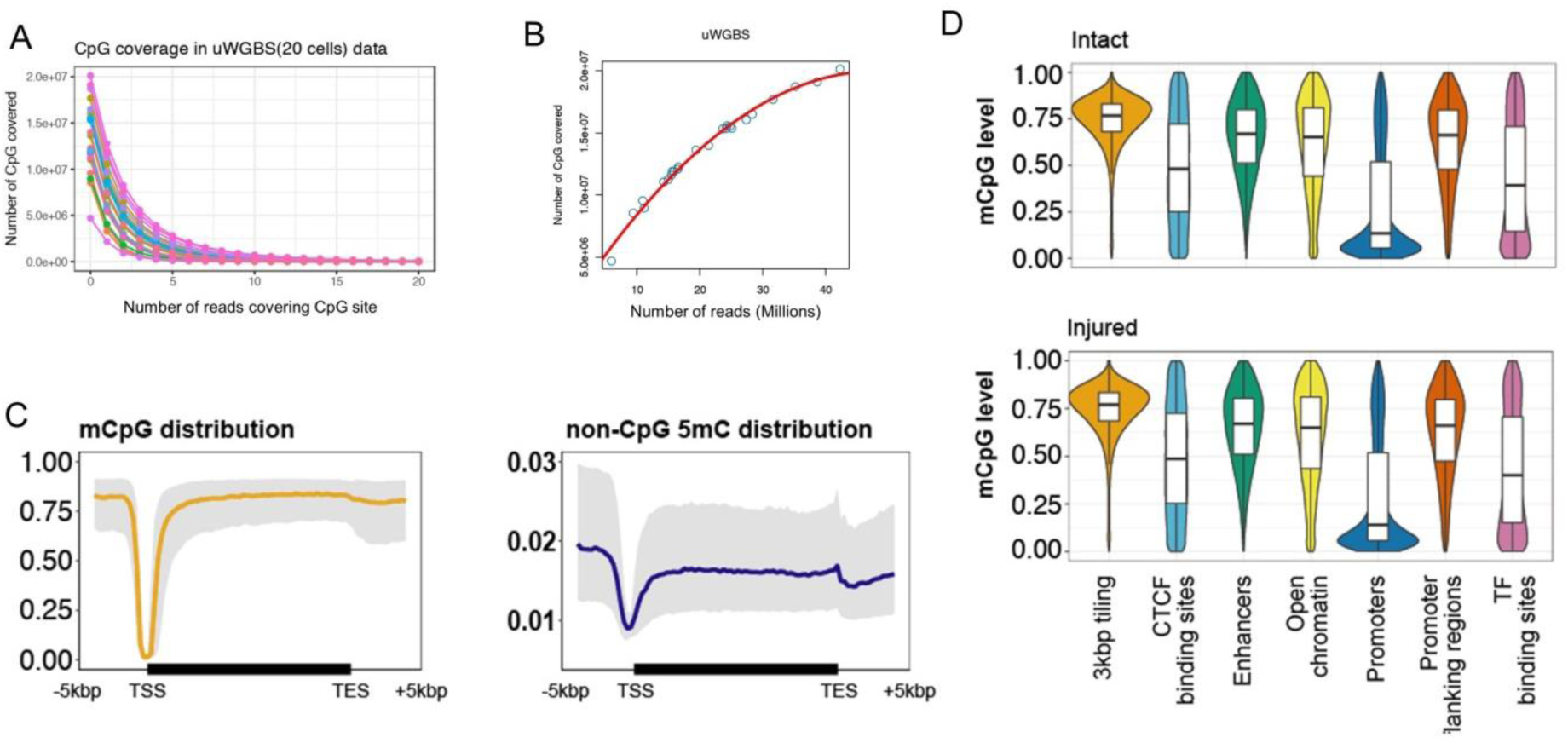
DNA methylation pattern of μWGBS. **A:** The number of CpG sites detected in each μWGBS data as a function of the number of reads covering the same CpG site. **B:** Relationship between number of reads in each library and number of CpG sites covered. **C:** Distribution of CpG and non-CpG methylation across gene bodies. For both CpG methylation and non-CpG methylation, we detected a hypomethylation valley around TSS regions. **D:** Distribution of CpG DNA methylation in different genomic contexts. The distribution of methylation level is very similar to what was previously reported by Farlik et.al (6).

**Fig. S5.**
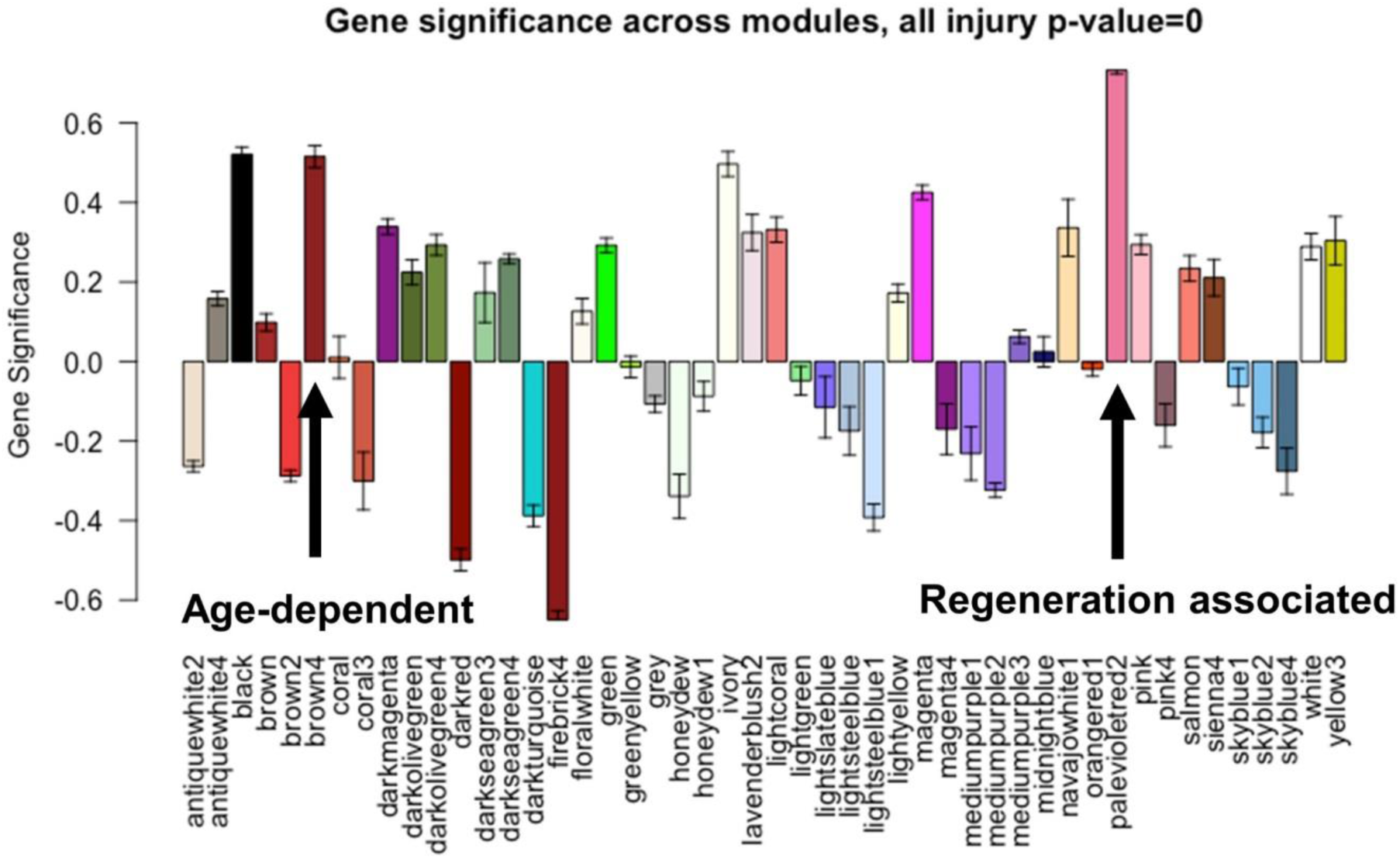
Module-trait correlations (Gene Significance) between 48 WGCNA modules and injury condition. The regeneration associated module and the age-dependent module have the highest correlation with injury condition.

**Fig S6.**
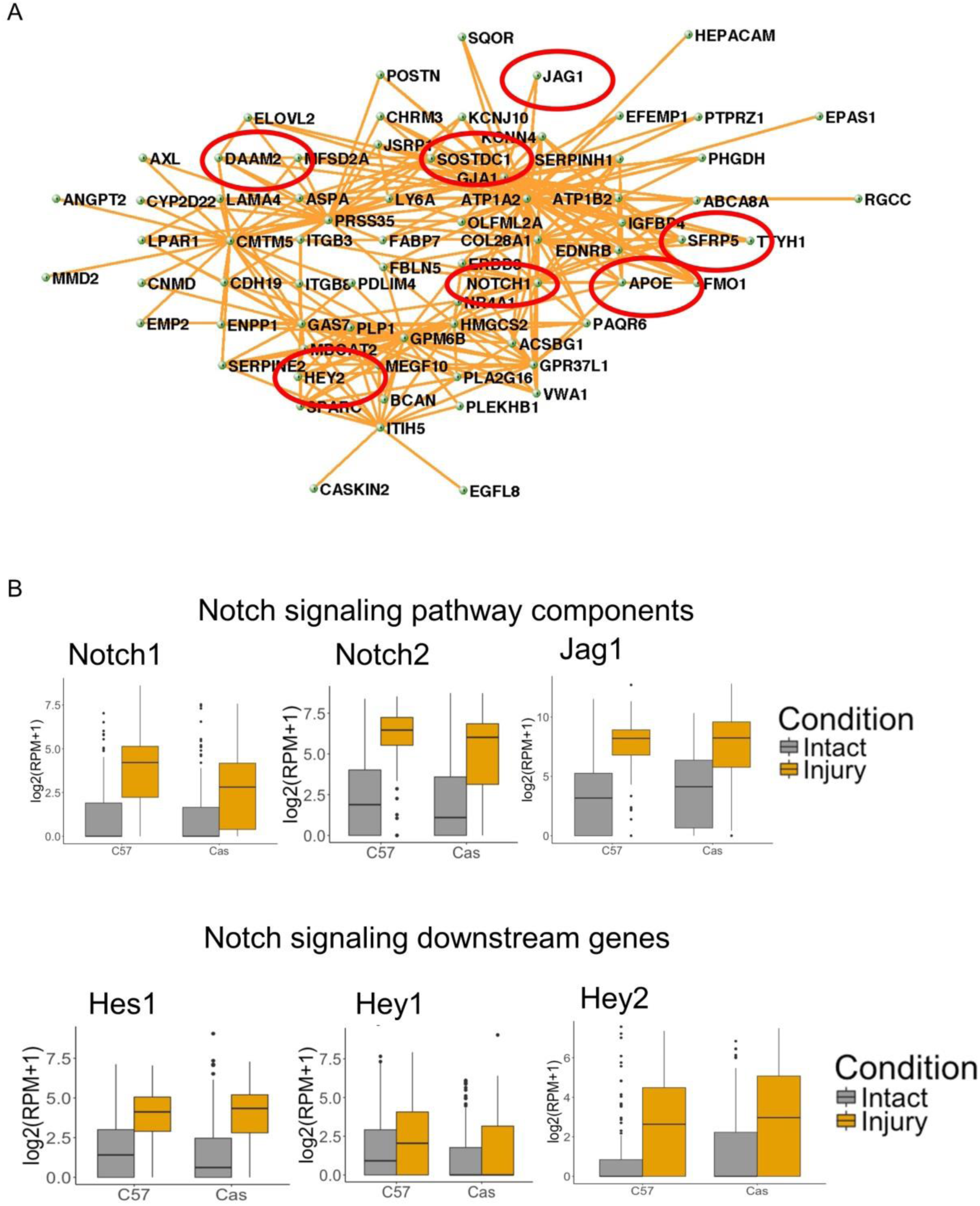
Notch signaling genes enriched in hub genes of RAG-module, and are upregulated by injury. **A:** The hub genes within the age-dependent module. Each node represents a gene, and the edge between two genes represents that these two genes are co-expressed. The genes related to the Notch signaling pathway are highlighted by red circles. **B:** Expression of Notch signaling components in DRG neurons of pure C57BL/6 and CAST/Ei mice, using data from Lisi, Veronique, et al (7). Notch signaling pathway components were upregulated by injury in DRG neurons.

**Fig. S7.**
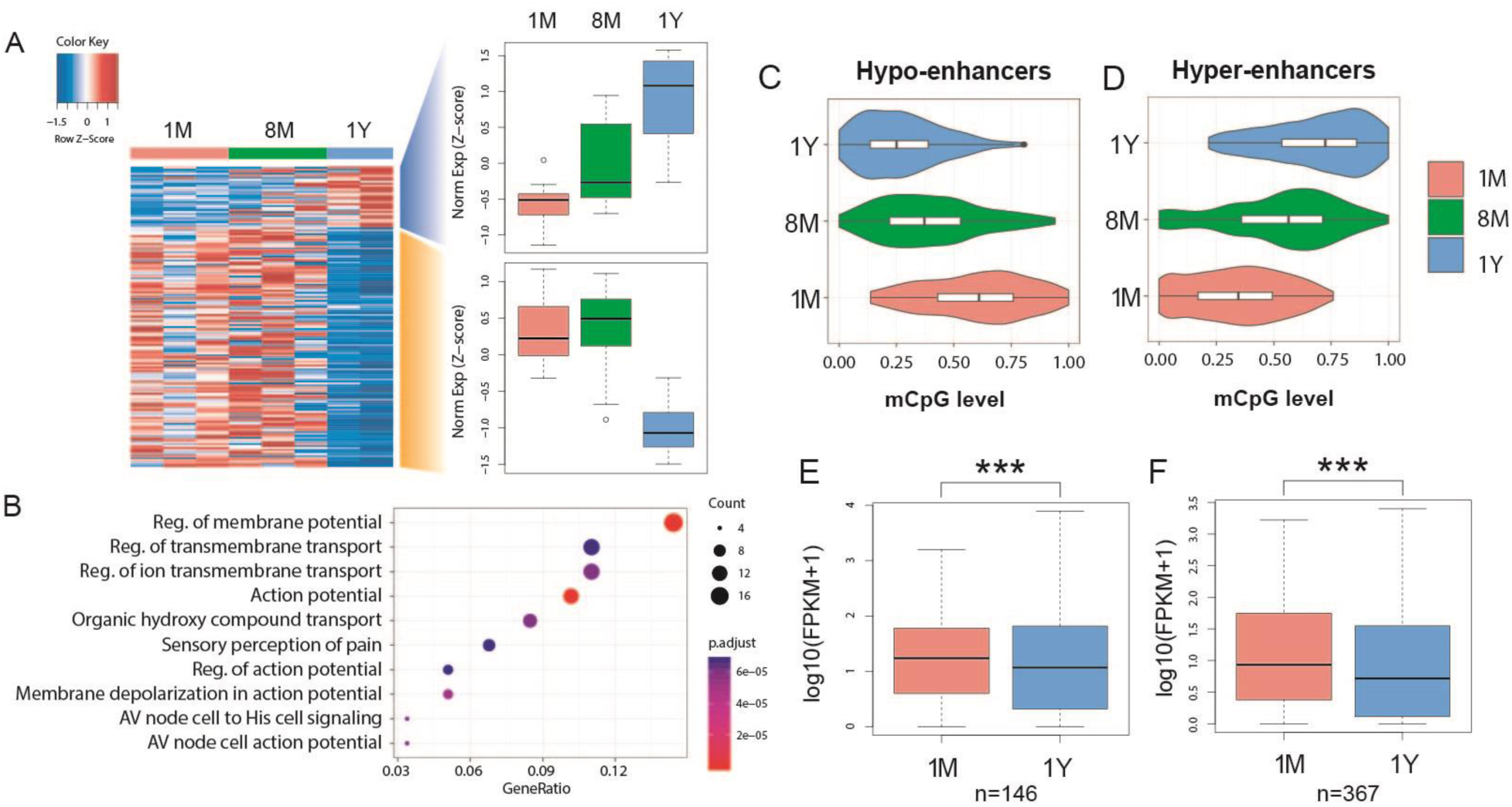
Age-dependent DNA methylation changes on enhancers are associated with age-dependent gene expression change. **A:** Differential expression analysis using µRNA-seq samples at different ages identified genes that are downregulated (yellow shade) and upregulated (blue shade) by age in mouse DRG neurons. The global expressions trend for upregulated and downregulated genes are shown as a boxplot. **B:** Gene ontology analysis showing the functional enrichment of age-downregulated genes. **C & D:** 254 enhancers were hypomethylated and 649 enhancers were hypermethylated by age between 1-month and 1-year neurons. **E & F:** Expression of target genes of differentially methylated enhancers were significantly downregulated, in spite of the direction of methylation change.

**Fig. S8.**
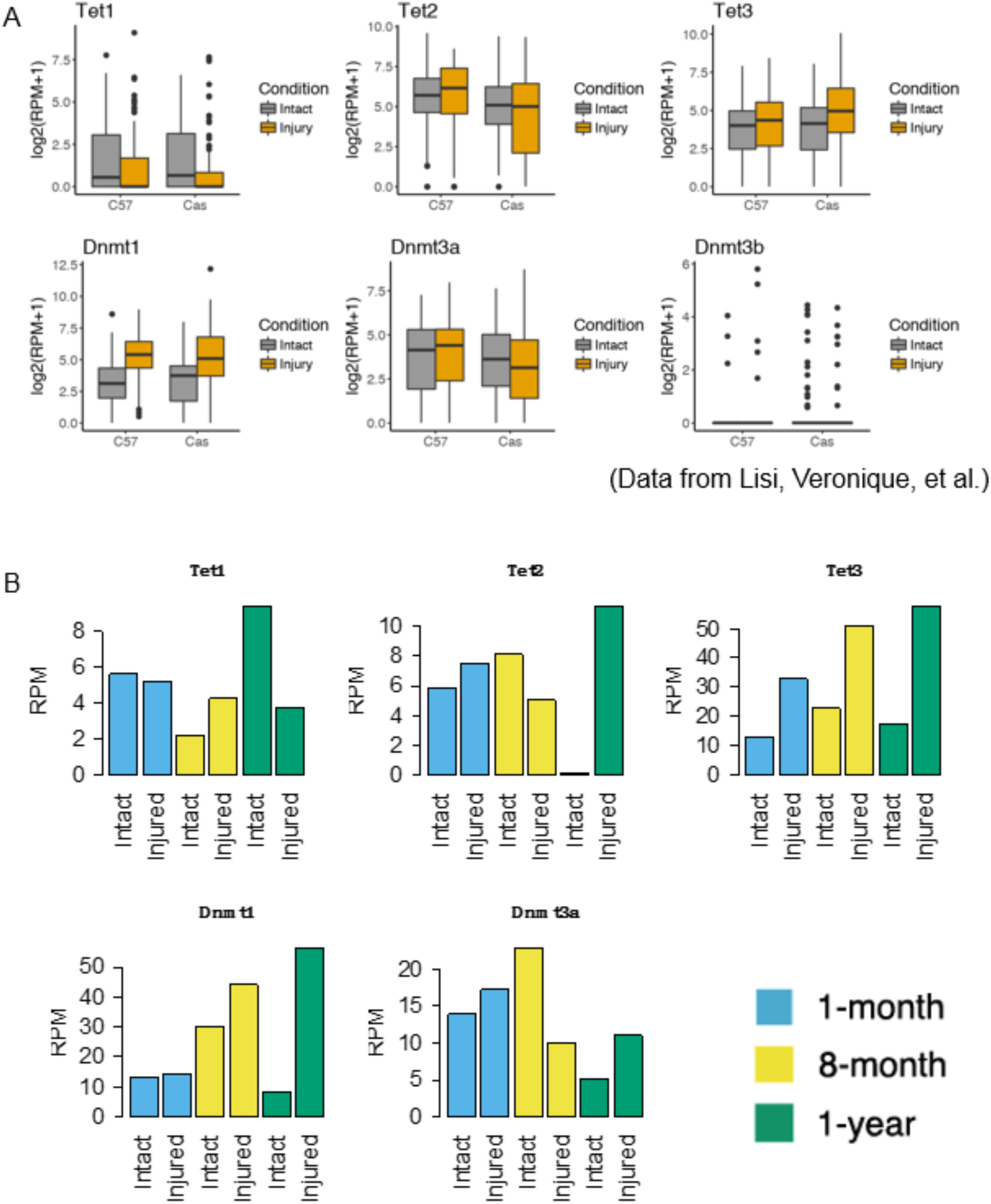
Expression change of DNA methylation related enzymes. **A:** Expression of DNMTs and TETs in intact and injured DRG neurons from C57BL/6 and CAST/Ei strain. Tet3 and Dnmt1 were consistently upregulated by injury in both C57BL/6 and CAST/Ei. **B:** Expression of DNMTs and TETs in intact and injured DRG neurons collected from our F1 hybrid mice at different age. Tet3 and Dnmt1 were consistently upregulated by injury, and their expression level in injured neurons are higher in aged neurons compared to young neurons.

**Fig S9.**
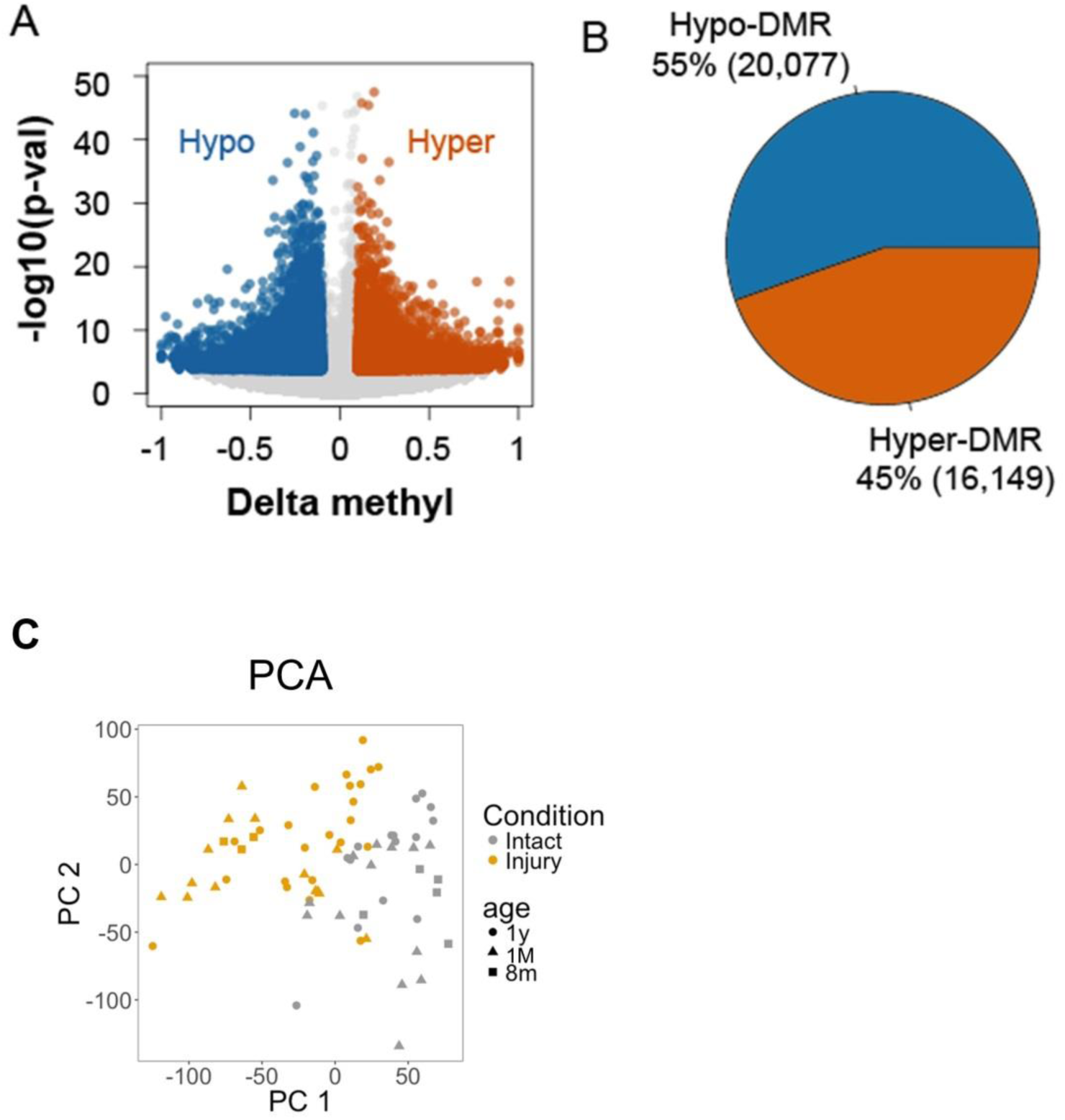
Differentially methylated regions (DMR) between injured and intact meta-DNA methylome. **A:** Volcano plot showing the methylation change and significance of all injury-induced DMRs. **B:** Number of hyper and hypo injury-induced DMRs. These DMRs were identified by combining data from all injured or intact samples into two meta-DNA methylomes, and identified the DMRs between them. These DMRs are collectively named as injury-induced DMRs. **C:** Visualization of µWGBS and scWGBS data principle component analysis.

**Fig. S10.**
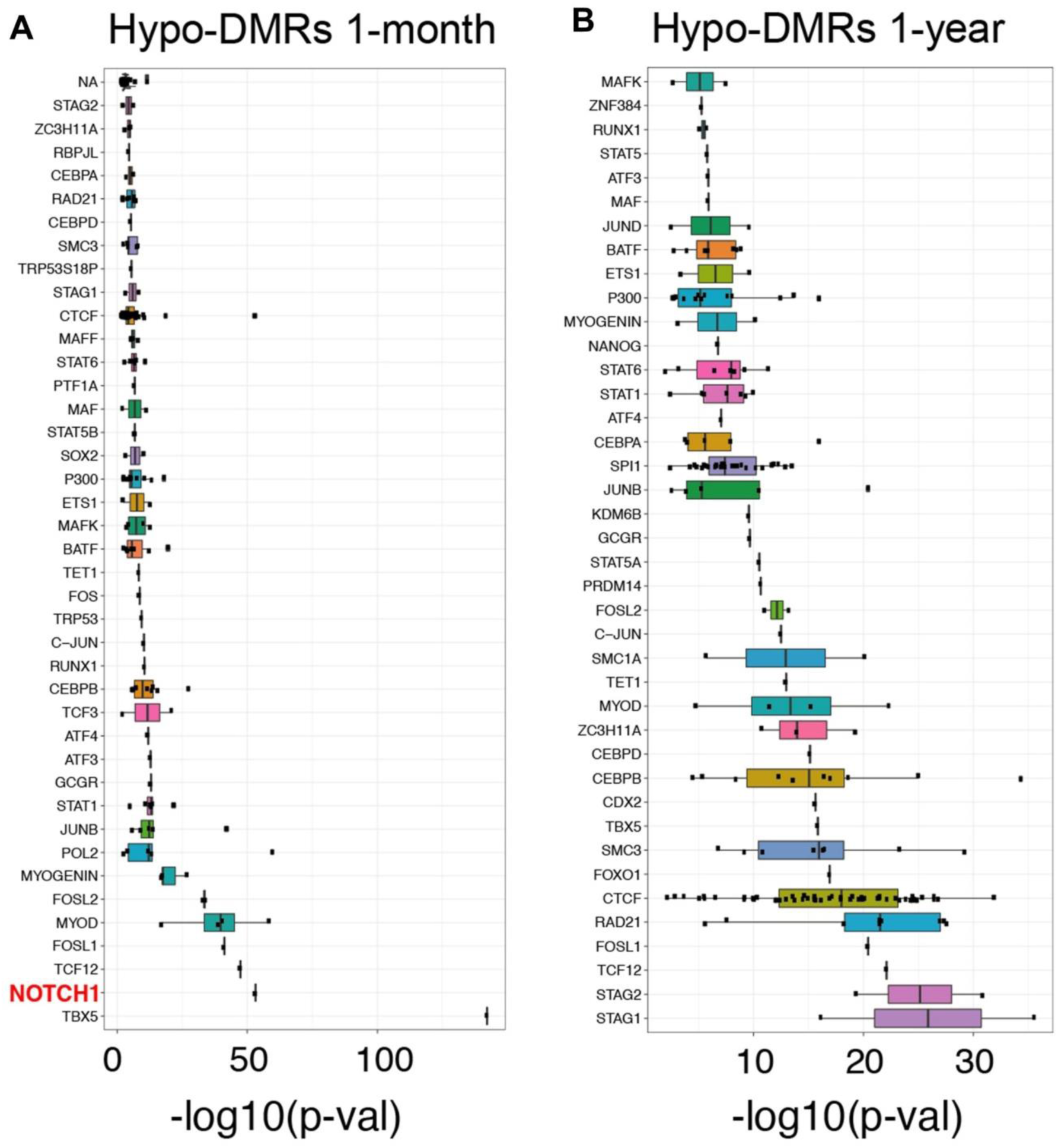
Boxplot showing the significance of the association between the hypo-methylated injury-induced DMRs and TF binding sites in 1-month and 1-year neurons, using LOLA. **A:** Boxplot showing the significance of the association between the hypo-methylated injury-induced DMRs and TF binding sites in 1-month neurons, using LOLA. **B:** Boxplot showing the significance of the association between the hypo-methylated injury-induced DMRs and TF binding sites in 1-year neurons. Notch1 binding sites were significantly overlapped with injury-caused hypo-DMRs in 1-month neurons, but not in 1-year neurons.

**Fig. S11.**
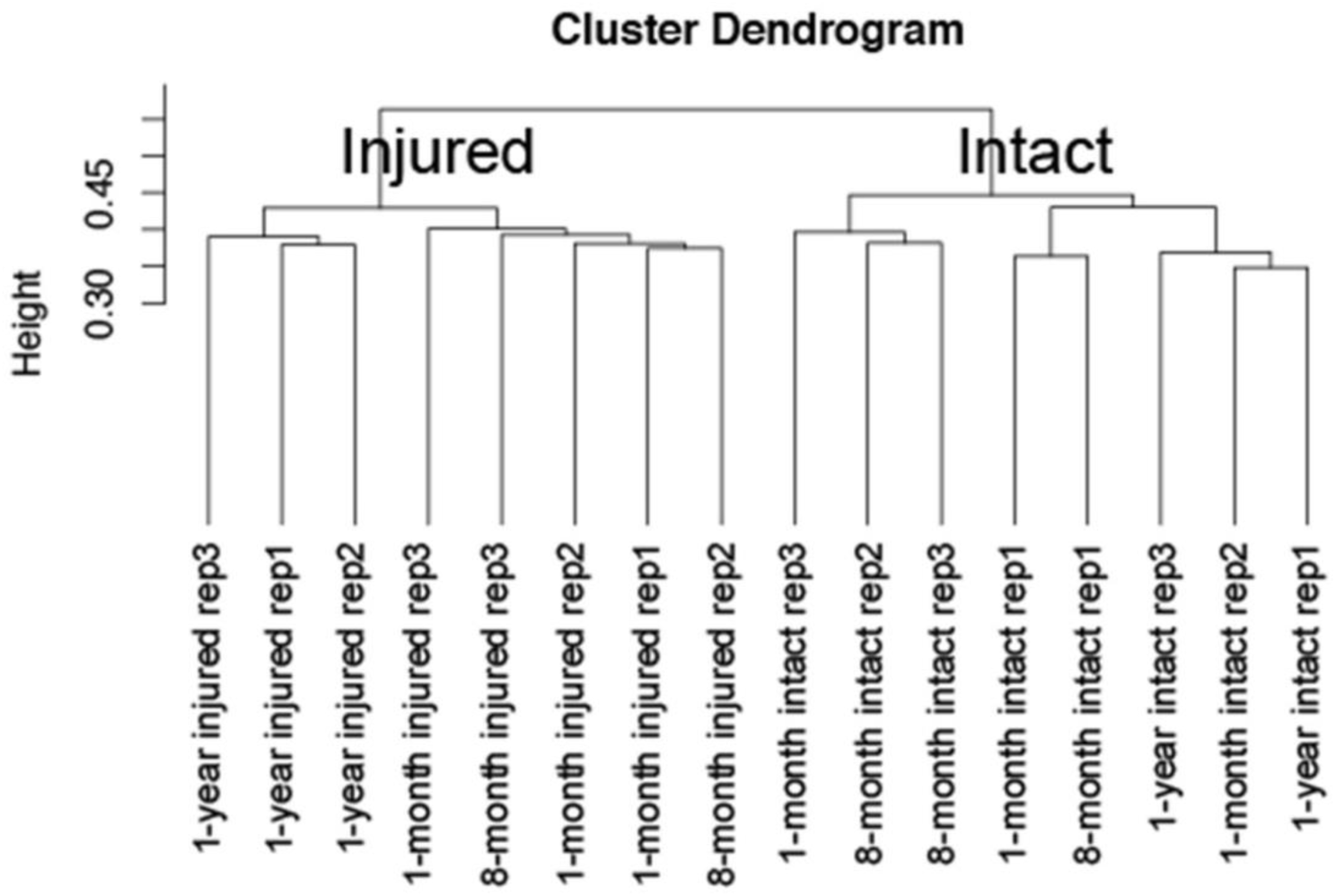
Hierarchical clustering of 16 μRNA-seq samples using all expressed 13,261 genes. Injured and intact samples were clearly separated.

**Fig. S12.**
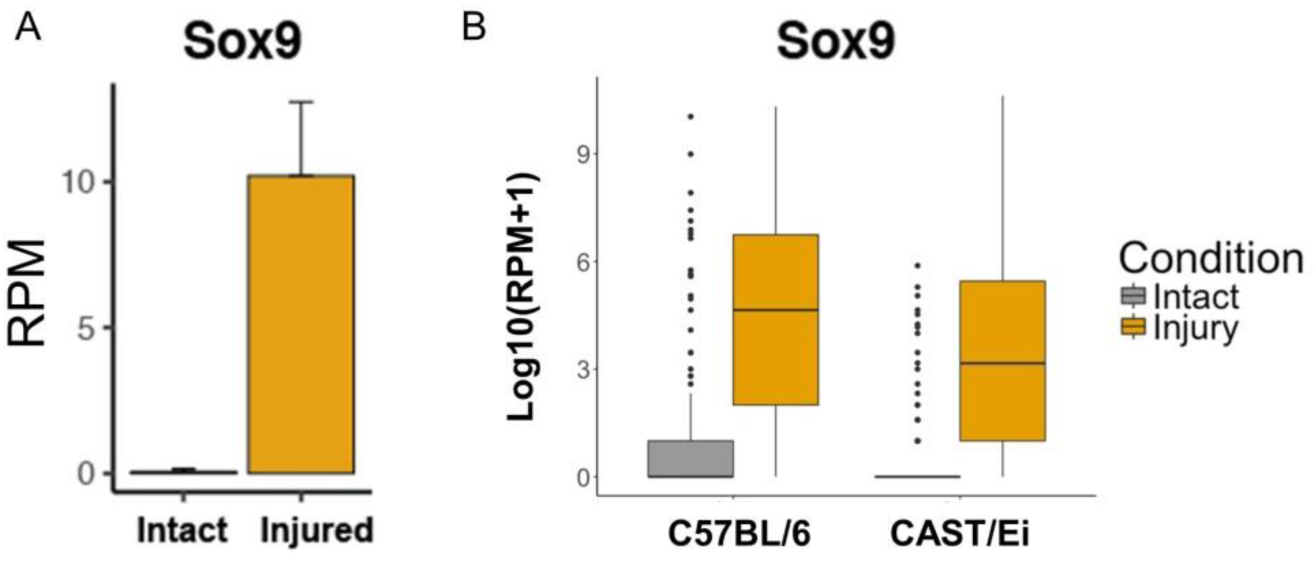
Expression of Sox9 in the μRNA-seq data generated in this study (A), and in a public single cell RNA-seq dataset from Lisi et al (7). (B). Sox9 was consistently upregulated after injury in both datasets.

**Fig. S13.**
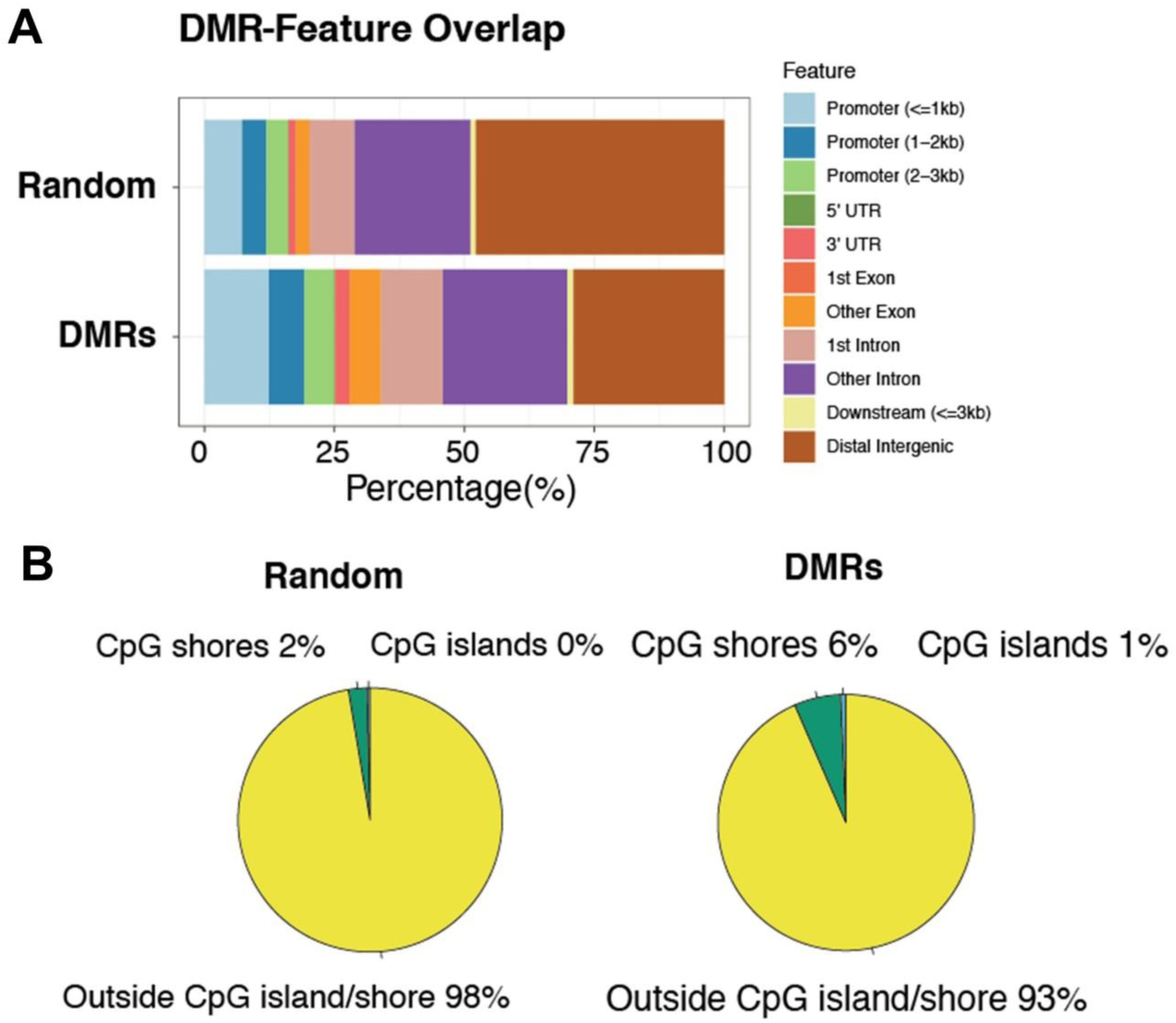
Enrichment of injury-induced DMRs. **A:** DMRs caused by injury are enriched in promoters and promoter flanking regions compared to random selected genomic regions with the same number and length. **B:** DMRs caused by injury are enriched in CpG islands and CpG shores compared to random selected genomic regions with the same number and length.

**Fig. S14.**
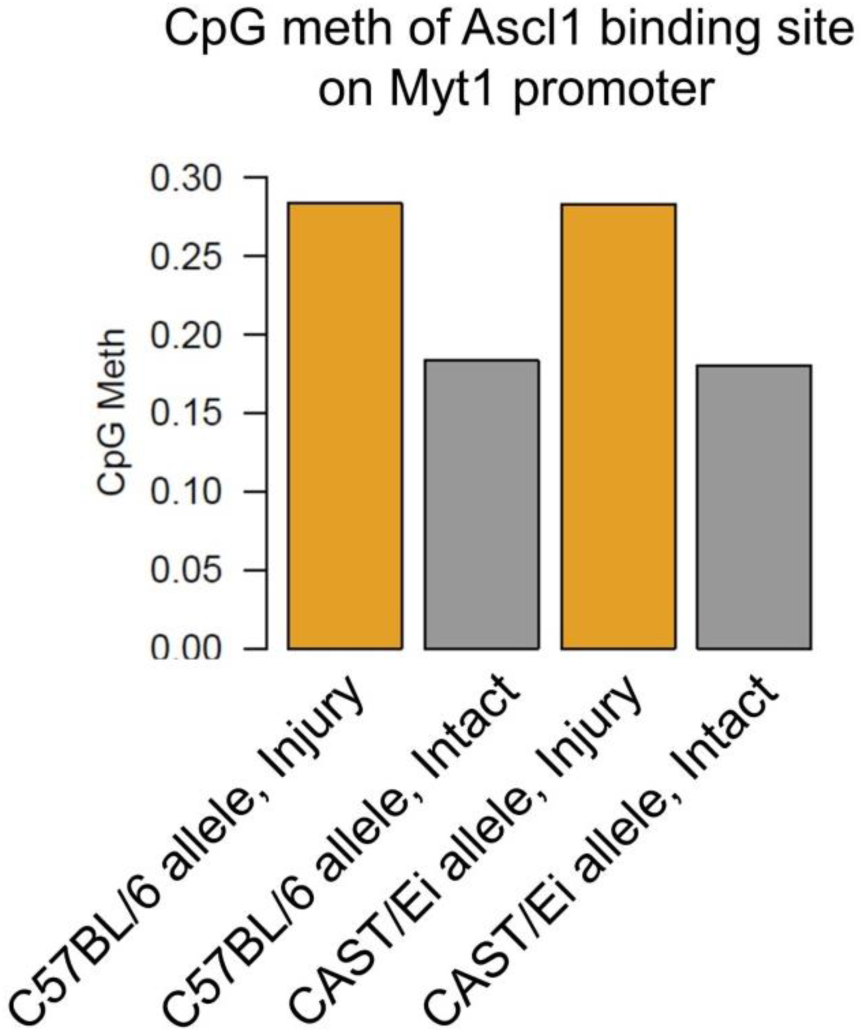
DNA methylation level of the Ascl1 binding site on the Myt1 promoter that is shown in Fig. 2B. The Ascl1 binding site on both C57BL/6 allele and CAST/Ei allele are hypermethylated after injury to the similar extent.

**Fig. S15.**
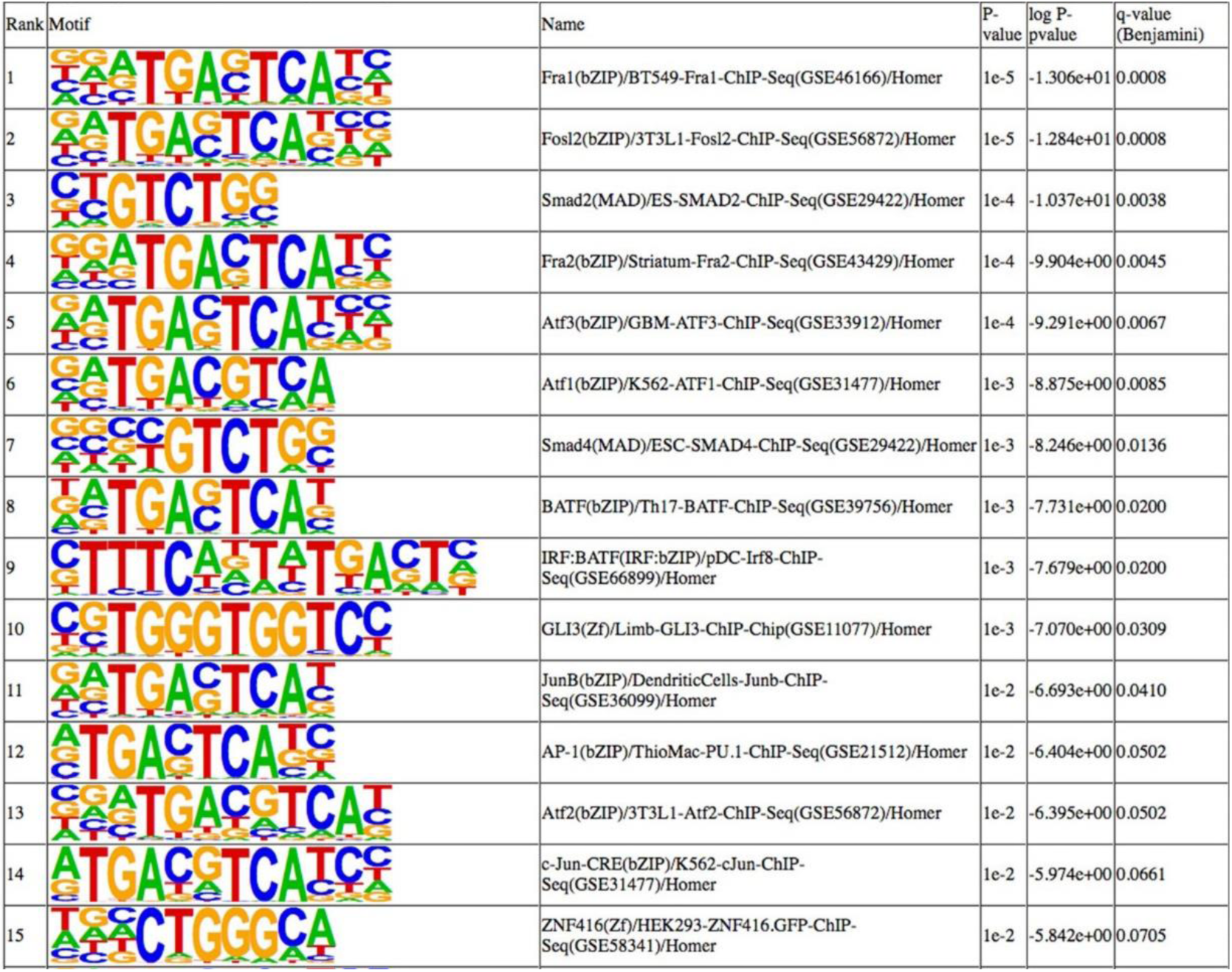
Complete list of transcription factor binding motifs enriched in promoters of genes in the regeneration associated WGCNA module.

**Fig. S16.**
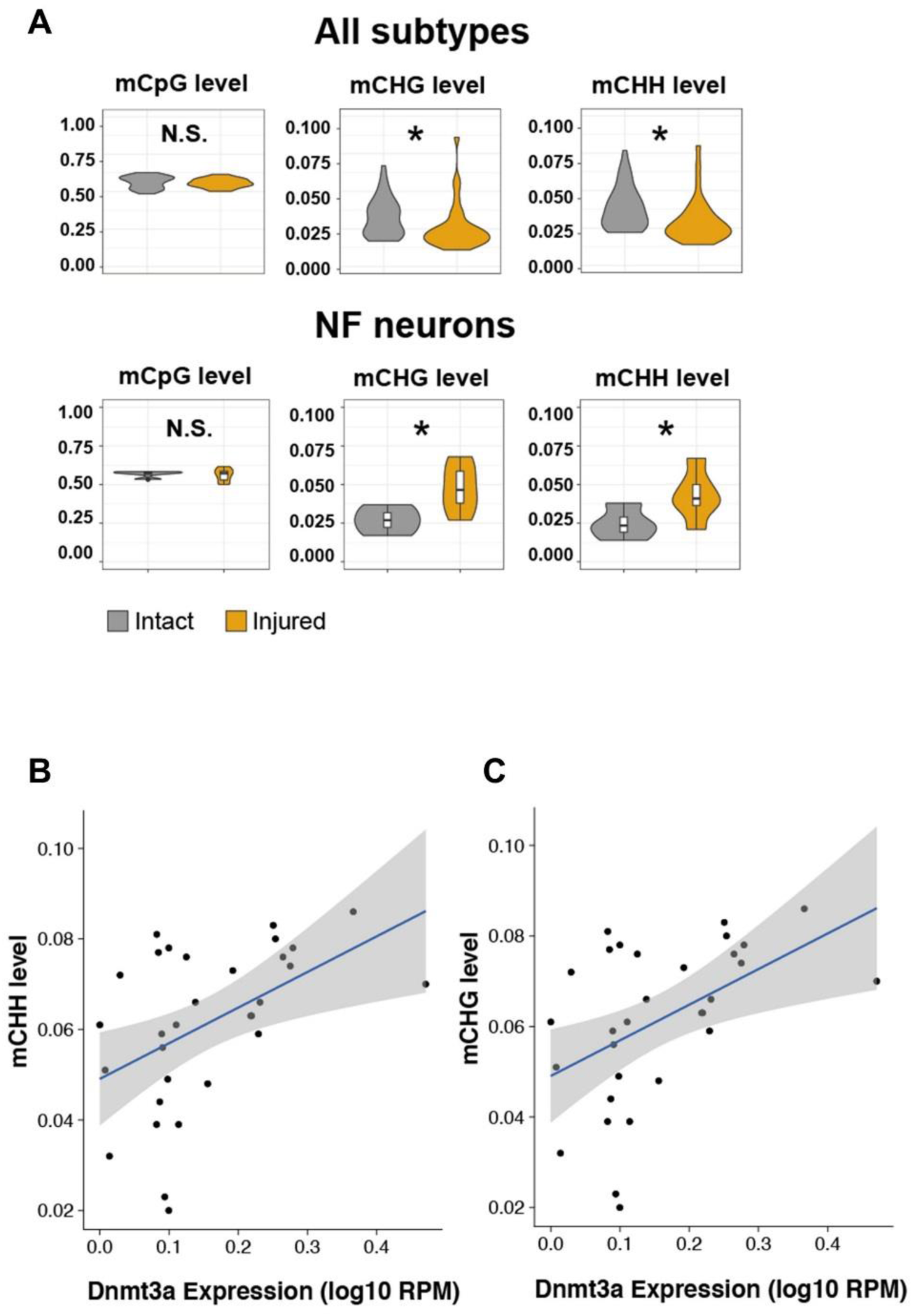
DNA methylation change in different neuron subtypes and correlation between DNA methylation level and Dnmt3a. **A:** Global CpG methylation and non-CpG methylation level between intact and injured WGBS samples. Top panel shown all neurons; bottom panel shows NF neurons. **B & C:** Non-CpG methylation is significantly correlated with Dnmt3a expression level from the same cell.

**Fig. S17.**
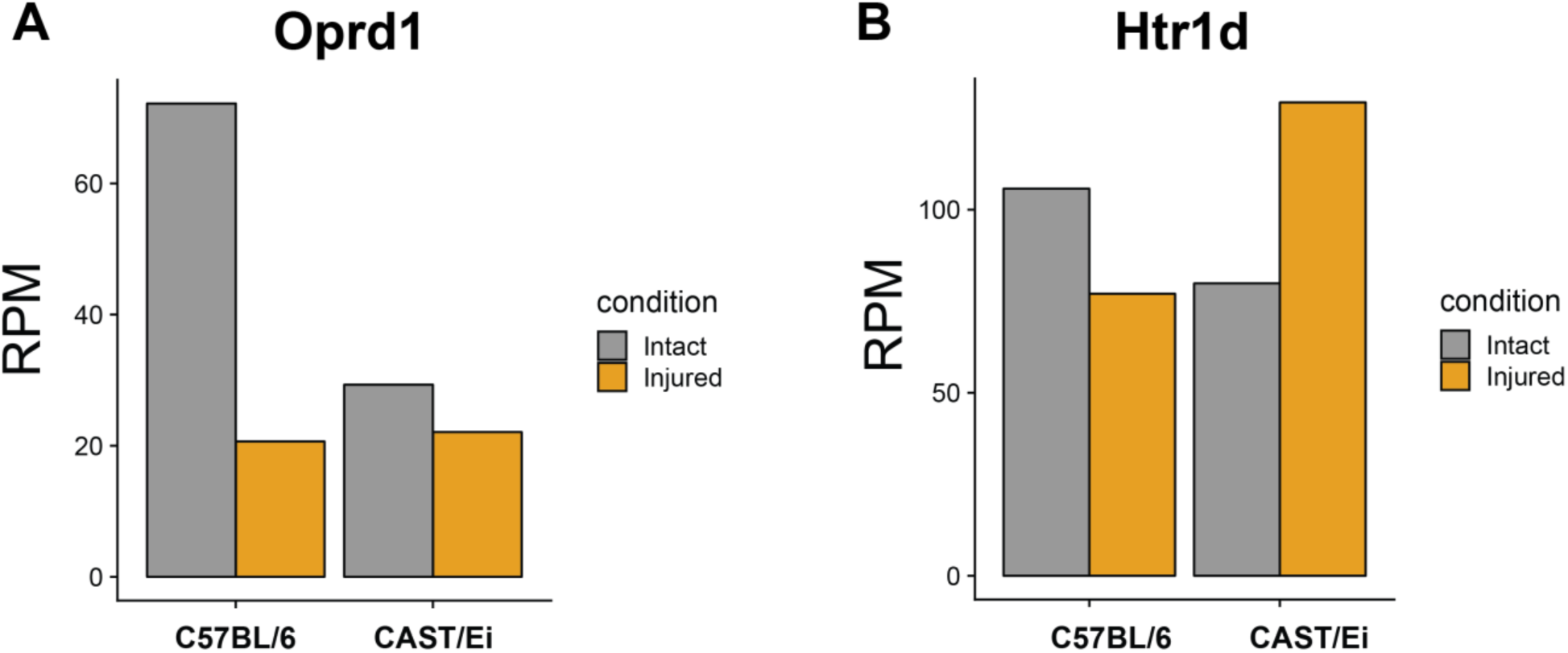
Expression change of representative cis-regulated genes after injury. **A:** Oprd1 expression from C57BL/6 allele is significantly downregulated after injury, but its expression from CAST/Ei allele downregulated with minor fold-change (Fisher’s exact test, BH corrected p-value = 2.14E-52). Consistently, Oprd1 promoter on C57BL/6 allele is significantly hypermethylated after injury, whereas Oprd1 promoter on CAST/Ei allele do not change significantly after injury. **B:** Htr1d expression from C57BL/6 allele is downregulated after injury, but its expression from CAST/Ei allele upregulated (Fisher’s exact test, BH corrected p-value = 1.28E-123). Consistently, Htr1d promoter on C57BL/6 allele is hypermethylated after injury, whereas Htr1d promoter on Cast/Ei allele is significantly hypomethylated.

**Fig S18.**
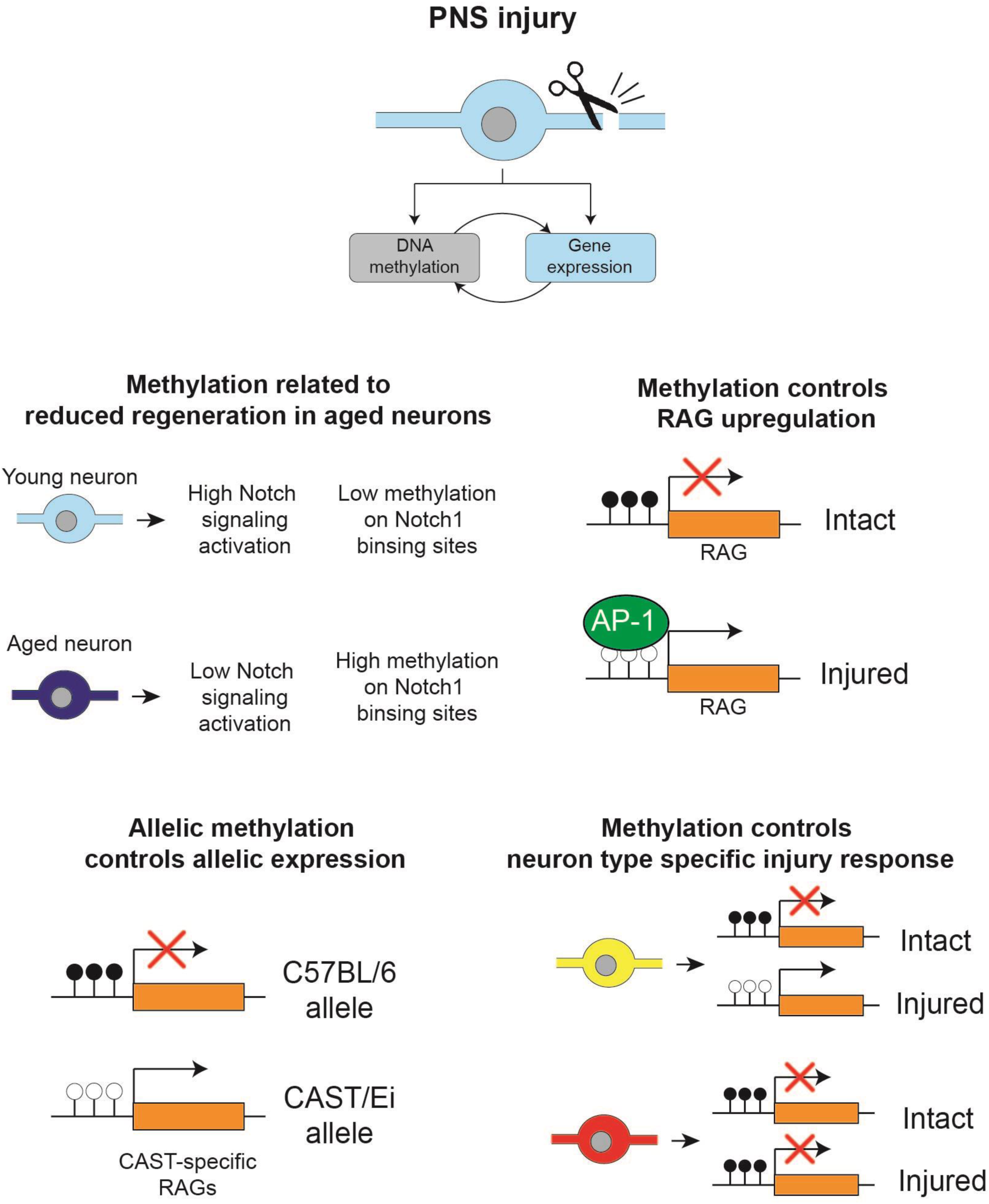
Schematic abstract summarizing four major points of the paper.

**Table S1:** Sample number after quality control.

| Library type | 1-month |  | 8-month |  | 1-year |  | SUM |
| --- | --- | --- | --- | --- | --- | --- | --- |
|  | intact | injured | intact | injured | intact | injured |  |
| μRNA-seq | 3 | 3 | 3 | 2 | 2 | 3 | 16 |
| μWGBS | 4 | 4 | 5 | 3 | 4 | 4 | 24 |
| scMT-seq (only RNA) | 7 | 7 | 0 | 0 | 0 | 6 | 20 |
| scMT-seq (only meth) | 2 | 3 | 0 | 0 | 4 | 5 | 14 |
| scMT-seq (dual-omic) | 7 | 7 | 0 | 0 | 7 | 14 | 35 |

**Table S2:**
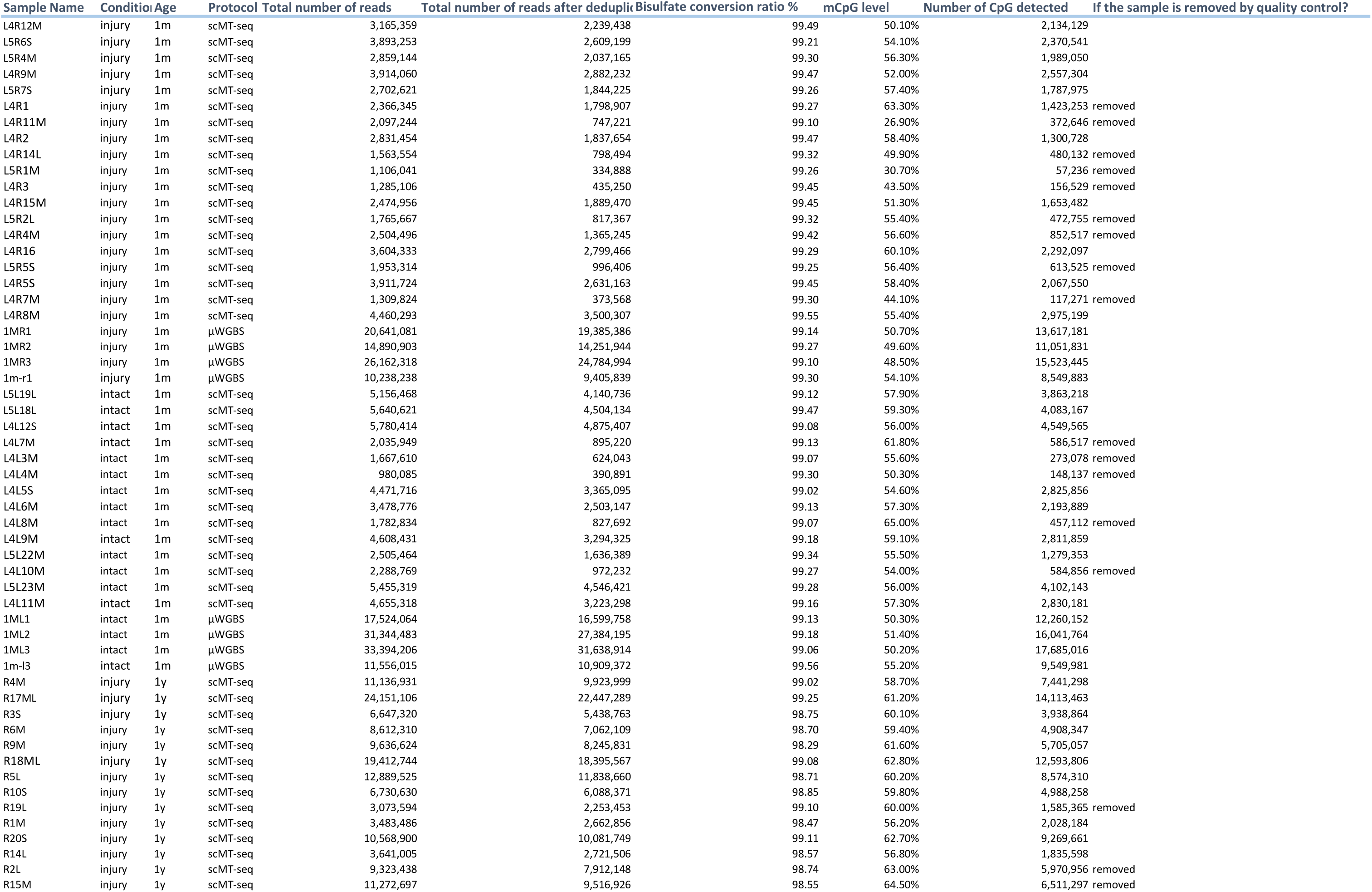
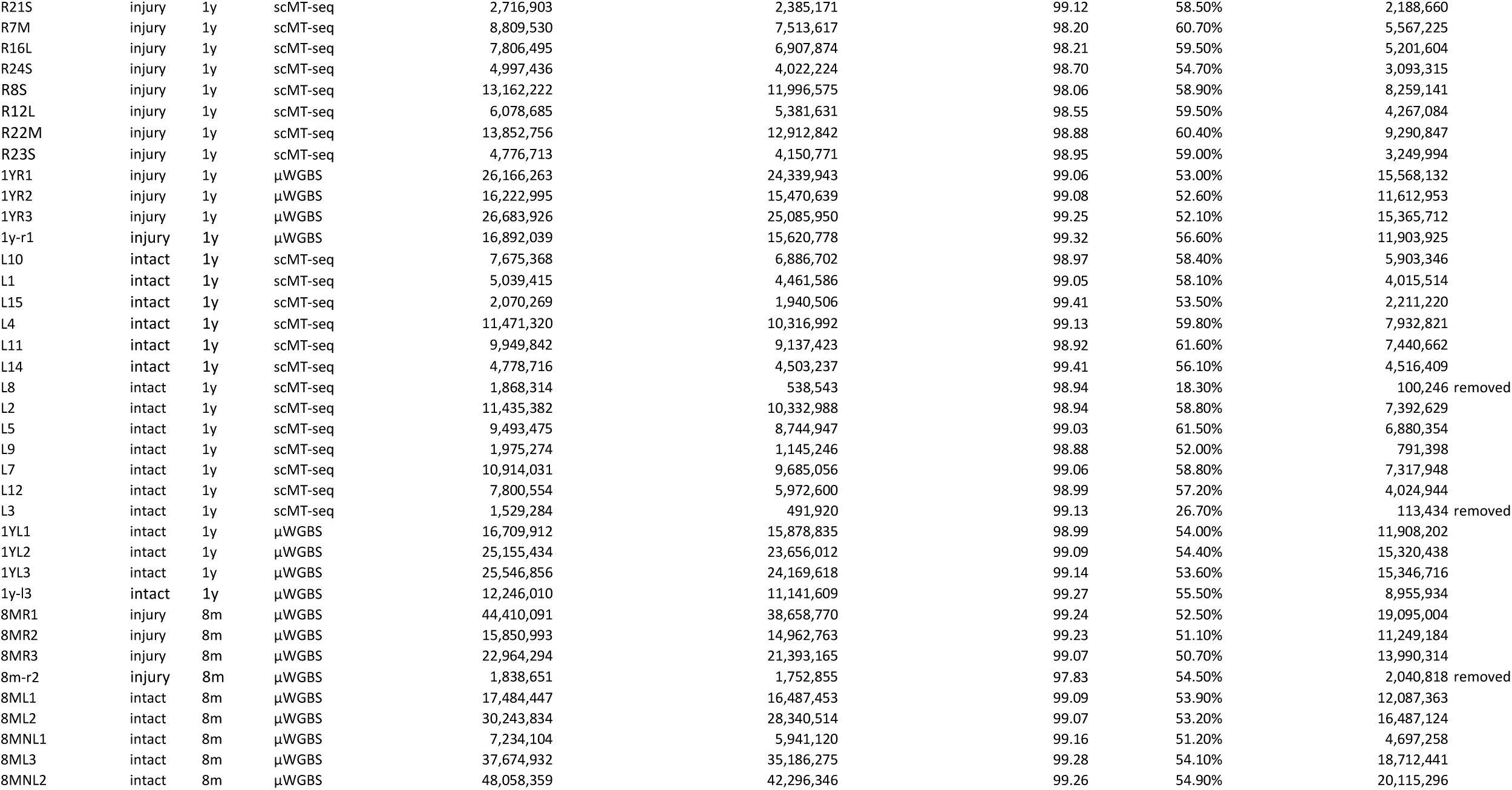
quality control table for DNA methylation data.

**Table S3:** quality control for RNA-seq data.

| Sample Name | Age | Condition | Protocol | Total reads | Uniquely mapped reads assigned to genes |
| --- | --- | --- | --- | --- | --- |
| 1m-l1 | 1m | intact | μRNA-seq | 26,988,277 | 11,745,664 |
| 1m-l2 | 1m | intact | μRNA-seq | 38,859,414 | 19,839,846 |
| 1ML3 | 1m | intact | μRNA-seq | 4,474,382 | 2,334,827 |
| 1m-l3 | 1m | intact | μRNA-seq | 27,394,678 | 12,819,962 |
| 1MR1 | 1m | injury | μRNA-seq | 3,299,638 | 1,646,489 |
| 1m-r1 | 1m | injury | μRNA-seq | 19,367,473 | 10,235,916 |
| 1m-r2 | 1m | injury | μRNA-seq | 18,605,757 | 10,026,828 |
| 1m-r3 | 1m | injury | μRNA-seq | 14,369,640 | 7,640,226 |
| 1y-l1 | 1y | intact | μRNA-seq | 42,479,341 | 15,837,615 |
| 1y-l2 | 1y | intact | μRNA-seq | 58,379,774 | 14,726,283 |
| 1YL3 | 1y | intact | μRNA-seq | 1,066,941 | 345,272 |
| 1y-l3 | 1y | intact | μRNA-seq | 33,217,115 | 12,127,372 |
| 1YR1 | 1y | injury | μRNA-seq | 6,078,415 | 3,169,052 |
| 1y-r1 | 1y | injury | μRNA-seq | 20,930,923 | 9,638,770 |
| 1y-r2 | 1y | injury | μRNA-seq | 23,261,846 | 11,651,079 |
| 1y-r3 | 1y | injury | μRNA-seq | 14,314,903 | 7,667,754 |
| 8m-l1 | 8m | intact | μRNA-seq | 28,850,281 | 15,989,989 |
| 8m-l2 | 8m | intact | μRNA-seq | 35,824,516 | 12,848,037 |
| 8m-l3 | 8m | intact | μRNA-seq | 35,575,795 | 16,216,660 |
| 8m-r1 | 8m | injury | μRNA-seq | 26,288,835 | 12,174,759 |
| 8m-r2 | 8m | injury | μRNA-seq | 22,220,570 | 11,324,380 |
| 8m-r3 | 8m | injury | μRNA-seq | 19,239,033 | 10,155,943 |
| L1 | 1y | intact | scMT-seq | 35,167,369 | 15,934,842 |
| L10 | 1y | intact | scMT-seq | 35,275,942 | 16,688,606 |
| L11 | 1y | intact | scMT-seq | 35,604,038 | 16,387,905 |
| L14 | 1y | intact | scMT-seq | 36,313,908 | 14,513,868 |
| L15 | 1y | intact | scMT-seq | 41,920,247 | 19,031,394 |
| L16 | 1y | intact | scMT-seq | 19,470,648 | 8,550,502 |
| L2 | 1y | intact | scMT-seq | 35,173,193 | 11,446,097 |
| L4 | 1y | intact | scMT-seq | 16,158,463 | 7,457,407 |
| L4L10M | 1m | intact | scMT-seq | 768,507 | 303,212 |
| L4L11M | 1m | intact | scMT-seq | 531,722 | 282,690 |
| L4L12S | 1m | intact | scMT-seq | 30,598,069 | 15,124,069 |
| L4L17S | 1m | intact | scMT-seq | 34,635,267 | 23,344,548 |
| L4L1M | 1m | intact | scMT-seq | 2,772,336 | 1,549,484 |
| L4L2L | 1m | intact | scMT-seq | 3,412,358 | 1,572,557 |
| L4L3M | 1m | intact | scMT-seq | 30,186,245 | 29,992,727 |
| L4L4M | 1m | intact | scMT-seq | 38,601,845 | 37,619,708 |
| L4L5S | 1m | intact | scMT-seq | 3,061,548 | 1,334,959 |
| L4L7M | 1m | intact | scMT-seq | 11,396,831 | 5,725,632 |
| L4L8M | 1m | intact | scMT-seq | 27,401,764 | 24,250,533 |
| L4L9M | 1m | intact | scMT-seq | 259,066 | 158,353 |
| L4R1 | 1m | injury | scMT-seq | 8,008,364 | 7,860,968 |
| L4R10S | 1m | intact | scMT-seq | 777,953 | 324,018 |
| L4R11M | 1m | intact | scMT-seq | 17,699,355 | 17,512,005 |
| L4R12m | 1m | injury | scMT-seq | 12,557,301 | 7,216,327 |
| L4R12M_2 | 1m | injury | scMT-seq | 5,625,280 | 3,342,289 |
| L4R13S | 1m | injury | scMT-seq | 878,865 | 425,350 |
| L4R14L | 1m | injury | scMT-seq | 23,803,863 | 12,198,374 |
| L4R15M | 1m | injury | scMT-seq | 18,083,421 | 16,581,944 |
| L4R16 | 1m | injury | scMT-seq | 17,614,489 | 9,123,441 |
| L4R2 | 1m | injury | scMT-seq | 3,732,219 | 1,642,304 |
| L4R4M | 1m | injury | scMT-seq | 2,796,749 | 1,364,904 |
| L4R6L | 1m | injury | scMT-seq | 3,174,611 | 1,567,178 |
| L4R8M | 1m | injury | scMT-seq | 25,153,768 | 24,559,636 |
| L4R9M | 1m | injury | scMT-seq | 23,770,528 | 11,009,186 |
| L5L18L | 1m | intact | scMT-seq | 15,291,125 | 9,442,406 |
| L5L19L | 1m | intact | scMT-seq | 20,302,011 | 10,537,098 |
| L5L20L | 1m | intact | scMT-seq | 111,683 | 50,830 |
| L5L21M | 1m | intact | scMT-seq | 37,903,327 | 37,688,607 |
| L5L22m | 1m | intact | scMT-seq | 30,777,806 | 15,893,493 |
| L5L22M_2 | 1m | intact | scMT-seq | 4,687,992 | 2,466,263 |
| L5L23M | 1m | intact | scMT-seq | 129,588 | 65,676 |
| L5L23M_2 | 1m | intact | scMT-seq | 2,505,200 | 1,201,377 |
| L5R1M | 1m | injury | scMT-seq | 14,482,203 | 14,404,431 |
| L5R2L | 1m | injury | scMT-seq | 35,450,092 | 12,926,813 |
| L5R3L | 1m | injury | scMT-seq | 10,517,354 | 5,820,075 |
| L5R4M | 1m | injury | scMT-seq | 14,883,237 | 8,778,012 |
| L5R5S | 1m | injury | scMT-seq | 29,741,221 | 18,259,732 |
| L5R6S | 1m | injury | scMT-seq | 11,815,259 | 6,644,461 |
| L5R7S | 1m | injury | scMT-seq | 12,502,987 | 6,797,808 |
| L5R8S | 1m | injury | scMT-seq | 19,377,096 | 12,567,423 |
| R10S | 1y | injury | scMT-seq | 31,705,395 | 10,194,737 |
| R12L | 1y | injury | scMT-seq | 24,888,666 | 7,110,067 |
| R13L | 1y | injury | scMT-seq | 22,839,140 | 6,611,336 |
| R14L | 1y | injury | scMT-seq | 22,016,337 | 6,312,472 |
| R14L_2 | 1y | injury | scMT-seq | 4,212,636 | 1,250,261 |
| R15M | 1y | injury | scMT-seq | 16,822,992 | 4,816,496 |
| R16L | 1y | injury | scMT-seq | 16,775,267 | 9,079,371 |
| R17ML | 1y | injury | scMT-seq | 40,628,655 | 26,743,967 |
| R18ML | 1y | injury | scMT-seq | 28,899,032 | 12,130,034 |
| R19L | 1y | injury | scMT-seq | 123,489 | 31,926 |
| R1M | 1y | injury | scMT-seq | 29,277,536 | 12,107,512 |
| R21S | 1y | injury | scMT-seq | 152,134 | 42,057 |
| R21S_2 | 1y | injury | scMT-seq | 2,401,417 | 1,003,018 |
| R23S | 1y | injury | scMT-seq | 3,345,884 | 1,133,819 |
| R24S | 1y | injury | scMT-seq | 25,843,028 | 11,335,654 |
| R2L | 1y | injury | scMT-seq | 28,572,301 | 12,650,581 |
| R3S | 1y | injury | scMT-seq | 34,238,710 | 12,573,918 |
| R4M | 1y | injury | scMT-seq | 42,046,361 | 19,417,116 |
| R5L | 1y | injury | scMT-seq | 25,128,852 | 7,384,743 |
| R6M | 1y | injury | scMT-seq | 18,464,723 | 6,061,828 |
| R6M_2 | 1y | injury | scMT-seq | 3,575,109 | 1,239,609 |
| R7M | 1y | injury | scMT-seq | 23,561,760 | 8,734,292 |
| R7M_2 | 1y | injury | scMT-seq | 4,654,621 | 2,137,547 |
| R8S | 1y | injury | scMT-seq | 25,506,842 | 8,871,680 |
| R9M | 1y | injury | scMT-seq | 14,328,407 | 4,938,187 |
| R9M_2 | 1y | injury | scMT-seq | 2,434,110 | 808,511 |

## References and Notes

1. Z. He, Y. Jin, Intrinsic Control of Axon Regeneration. Neuron 90, 437–451 (2016).

2. K. Liu, A. Tedeschi, K. K. Park, Z. G. He, Neuronal Intrinsic Mechanisms of Axon Regeneration. Annu Rev Neurosci 34, 131–152 (2011).

3. F. Rossi, S. Gianola, L. Corvetti, Regulation of intrinsic neuronal properties for axon growth and regeneration. Prog Neurobiol 81, 1–28 (2007).

4. F. Q. Zhou, W. D. Snider, Intracellular control of developmental and regenerative axon growth. Philos T R Soc B 361, 1575–1592 (2006).

5. J. Y. Zhao et al., DNA methyltransferase DNMT3a contributes to neuropathic pain by repressing Kcna2 in primary afferent neurons. Nature Communications 8, (2017).

6. Y. L. Weng et al., An Intrinsic Epigenetic Barrier for Functional Axon Regeneration. Neuron 94, 337-+ (2017).

7. M. Golzenleuchter et al., Plasticity of DNA methylation in a nerve injury model of pain. Epigenetics-Us 10, 200–212 (2015).

8. G. Laumet et al., G9a is essential for epigenetic silencing of K+ channel genes in acute- to-chronic pain transition. Nat Neurosci 18, 1746–1755 (2015).

9. G. L. Hu et al., Single-cell RNA-seq reveals distinct injury responses in different types of DRG sensory neurons. Sci Rep-Uk 6, (2016).

10. Y. Hu et al., Simultaneous profiling of transcriptome and DNA methylome from a single cell. Genome biology 17, 88 (2016).

11. Y. Hou et al., Single-cell triple omics sequencing reveals genetic, epigenetic, and transcriptomic heterogeneity in hepatocellular carcinomas. Cell Research 26, 304–319 (2016).

12. Z. G. Xue et al., Genetic programs in human and mouse early embryos revealed by single-cell RNA sequencing. Nature 500, 593-+ (2013).

13. P. Langfelder, S. Horvath, WGCNA: an R package for weighted correlation network analysis. Bmc Bioinformatics 9, (2008).

14. R. Lister et al., Global epigenomic reconfiguration during mammalian brain development. Science 341, 1237905 (2013).

15. A. D. King et al., Reversible Regulation of Promoter and Enhancer Histone Landscape by DNA Methylation in Mouse Embryonic Stem Cells. Cell Rep 17, 289–302 (2016).

16. N. C. Sheffield, C. Bock, LOLA: enrichment analysis for genomic region sets and regulatory elements in R and Bioconductor. Bioinformatics 32, 587–589 (2016).

17. C. Altmann et al., Progranulin promotes peripheral nerve regeneration and reinnervation: role of notch signaling. Mol Neurodegener 11, 69 (2016).

18. S. A. Bonini, G. Ferrari-Toninelli, M. Montinaro, M. Memo, Notch signalling in adult neurons: a potential target for microtubule stabilization. Ther Adv Neurol Diso 6, 375–385 (2013).

19. X. F. Ding, X. Gao, X. C. Ding, M. Fan, J. Chen, Postnatal dysregulation of Notch signal disrupts dendrite development of adult-born neurons in the hippocampus and contributes to memory impairment. Sci Rep 6, 25780 (2016).

20. J. Terragni, et al., Notch signaling genes: myogenic DNA hypomethylation and 5-hydroxymethylcytosine. Epigenetics-Us 9, 842–850 (2014).

21. R. Seijffers, C. D. Mills, C. J. Woolf, ATF3 increases the intrinsic growth state of DRG neurons to enhance peripheral nerve regeneration. J Neurosci 27, 7911–7920 (2007).

22. H. Tsujino et al., Activating transcription factor 3 (ATF3) induction by axotomy in sensory and motoneurons: A novel neuronal marker of nerve injury. Mol Cell Neurosci 15, 170–182 (2000).

23. G. Raivich et al., The AP-1 transcription factor c-jun is required for efficient axonal regeneration. Neuron 43, 57–67 (2004).

24. M. A. Riyadh, Y. Shinmyo, K. Ohta, H. Tanaka, Inhibitory effects of draxin on axonal outgrowth and migration of precerebellar neurons. Biochem Bioph Res Co 449, 169–174 (2014).

25. M. Robinson et al., FLRT3 is expressed in sensory neurons after peripheral nerve injury and regulates neurite outgrowth. Mol Cell Neurosci 27, 202–214 (2004).

26. Y. H. Wei et al., Nrf2 in ischemic neurons promotes retinal vascular regeneration through regulation of semaphorin 6A. P Natl Acad Sci USA 112, E6927–E6936 (2015).

27. M. Cheung, J. Briscoe, Neural crest development is regulated by the transcription factor Sox9. Development 130, 5681–5693 (2003).

28. W. M. McKillop et al., Conditional Sox9 ablation improves locomotor recovery after spinal cord injury by increasing reactive sprouting. Exp Neurol 283, 1–15 (2016).

29. H. Kanno et al., Genetic Ablation of Transcription Repressor Bach1 Reduces Neural Tissue Damage and Improves Locomotor Function after Spinal Cord Injury in Mice. J Neurotraum 26, 31–39 (2009).

30. M. Gey et al., Atf3 mutant mice show reduced axon regeneration and impaired regeneration-associated gene induction after peripheral nerve injury. Open Biol 6, (2016).

31. K. K. Rau et al., Cutaneous tissue damage induces long-lasting nociceptive sensitization and regulation of cellular stress- and nerve injury-associated genes in sensory neurons. Exp Neurol 283, 413–427 (2016).

32. T. Wang et al., Phenotypic Switching of Nonpeptidergic Cutaneous Sensory Neurons following Peripheral Nerve Injury. Plos One 6, (2011).

33. Y. H. Wang et al., Peripheral Nerve Injury Induces Down-Regulation of Foxo3a and p27(kip1) in Rat Dorsal Root Ganglia. Neurochem Res 34, 891–898 (2009).

34. F. F. Vasconcelos et al., MyT1 Counteracts the Neural Progenitor Program to Promote Vertebrate Neurogenesis. Cell Rep 17, 469–483 (2016).

35. H. D. MacGillavry et al., Genome-wide gene expression and promoter binding analysis identifies NFIL3 as a repressor of C/EBP target genes in neuronal outgrowth. Mol Cell Neurosci 46, 460–468 (2011).

36. H. D. MacGillavry et al., NFIL3 and cAMP Response Element-Binding Protein Form a Transcriptional Feedforward Loop that Controls Neuronal Regeneration-Associated Gene Expression. J Neurosci 29, 15542–15550 (2009).

37. C. Altmann et al., Progranulin promotes peripheral nerve regeneration and reinnervation: role of notch signaling. Molecular Neurodegeneration 11, (2016).

38. K. Ben-Yaakov et al., Axonal transcription factors signal retrogradely in lesioned peripheral nerve. Embo J 31, 1350–1363 (2012).

39. Y. Lu, S. Belin, Z. G. He, Signaling regulations of neuronal regenerative ability. Curr Opin Neurobiol 27, 135–142 (2014).

40. S. Nadeau, P. Hein, K. J. L. Fernandes, A. C. Peterson, F. D. Miller, A transcriptional role for C/EBP beta in the neuronal response to axonal injury. Mol Cell Neurosci 29, 525–535 (2005).

41. A. M. Kenney, J. D. Kocsis, Peripheral axotomy induces long-term c-Jun amino-terminal kinase-1 activation and activator protein-1 binding activity by c-Jun and junD in adult rat dorsal root ganglia in vivo. J Neurosci 18, 1318–1328 (1998).

42. T. Omura et al., Robust Axonal Regeneration Occurs in the Injured CAST/Ei Mouse CNS. Neuron 86, 1215–1227 (2015).

43. F. E. Holmes et al., Targeted disruption of the galanin gene reduces the number of sensory neurons and their regenerative capacity. Proc Natl Acad Sci U S A 97, 11563–11568 (2000).

44. L. Liang et al., G9a inhibits CREB-triggered expression of mu opioid receptor in primary sensory neurons following peripheral nerve injury. Mol Pain 12, (2016).

45. S. H. Avila-Rojas et al., Role of spinal 5-HT5A, and 5-HT1A/1B/1D, receptors in neuropathic pain induced by spinal nerve ligation in rats. Brain Res 1622, 377–385 (2015).

46. Q. L. Deng, D. Ramskold, B. Reinius, R. Sandberg, Single-Cell RNA-Seq Reveals Dynamic, Random Monoallelic Gene Expression in Mammalian Cells. Science 343, 193–196 (2014).

47. B. A. Chestnut et al., Epigenetic Regulation of Motor Neuron Cell Death through DNA Methylation. J Neurosci 31, 16619–16636 (2011).

48. Y. F. Li et al., Stella safeguards the oocyte methylome by preventing de novo methylation mediated by DNMT1. Nature 564, 136-+ (2018).

49. J. Feng et al., Dnmt1 and Dnmt3a maintain DNA methylation and regulate synaptic function in adult forebrain neurons. Nat Neurosci 13, 423 (2010).

## References

1. G. L. Hu et al., Single-cell RNA-seq reveals distinct injury responses in different types of DRG sensory neurons. Sci Rep-Uk 6, (2016).

2. Y. Hu et al., Simultaneous profiling of transcriptome and DNA methylome from a single cell. Genome biology 17, 88 (2016).

3. S. A. Malin, B. M. Davis, D. C. Molliver, Production of dissociated sensory neuron cultures and considerations for their use in studying neuronal function and plasticity. Nat Protoc 2, 152–160 (2007).

4. S. A. Smallwood et al., Single-cell genome-wide bisulfite sequencing for assessing epigenetic heterogeneity. Nat Methods 11, 817–820 (2014).

5. D. Usoskin et al., Unbiased classification of sensory neuron types by large-scale single-cell RNA sequencing. Nat Neurosci 18, 145 (2015).

6. M. Farlik et al., DNA Methylation Dynamics of Human Hematopoietic Stem Cell Differentiation. Cell Stem Cell 19, 808–822 (2016).

7. V. Lisi et al., Enhanced Neuronal Regeneration in the CAST/Ei Mouse Strain Is Linked to Expression of Differentiation Markers after Injury. Cell Rep 20, 1136–1147 (2017).

